# Embryonic macrophages chronologically change their brain-entry routes and competence for neural progenitor-induced microglial differentiation

**DOI:** 10.64898/2026.10.22.760592

**Authors:** Mizuki Ono, Tsukasa Shimamura, Ziwei Yin, Takaki Miyata, Yuki Hattori

## Abstract

Microglia arise from macrophage precursors that enter the developing brain and acquire microglial identity, but how these processes are coordinated remains unclear. We show that despite prominent entry of macrophages from the ventricles into the mouse telencephalon at embryonic day (E) 12.5, they enter from the meninges at E11.5, preferentially in the rostral pallium. Consistent with this regional pattern, whole-mount en face imaging revealed that focal holes in the pial basement membrane were most frequently observed in the rostral pallium. We found that neural progenitors, but not endothelial cells, promote acquisition of microglial properties through a process that requires close cellular proximity and partly depends on TGF-β receptor signaling. This responsiveness is substantially stronger in E13.5 than in E17.5 meningeal macrophages, indicating stage-dependent competence to acquire microglial properties. These findings suggest that microglial establishment is developmentally regulated by macrophage access to the telencephalon and responsiveness to local neural cues.

## Introduction

Microglia are the resident macrophages of the central nervous system (CNS) and play diverse roles in tissue homeostasis, immune surveillance, and neural development (Paolicelli et al., 2011, Parkhurst et al., 2013, Colonna and Butovsky, 2017, Depp et al., 2025). Fate-mapping studies have established that microglia arise predominantly from primitive macrophage progenitors generated in the yolk sac (Ginhoux et al., 2010, Kierdorf et al., 2013, Shimizu and Prinz, 2025). These progenitors colonize the developing CNS early in embryogenesis and progressively acquire the transcriptional and functional properties characteristic of microglia through a stepwise developmental program (Matcovitch-Natan et al., 2016, Li et al., 2019, Kracht et al., 2020, Bridlance et al., 2026). This differentiation is strongly influenced by signals from the local CNS environment, which contribute to the establishment of microglial identity (Gosselin et al., 2017, Bennett et al., 2018, Ayata et al., 2018). Thus, establishment of the microglial population depends on two fundamental processes: the entry of macrophage precursors into the developing brain and their subsequent differentiation within the neural environment.

The CNS microenvironment provides critical signals that promote the acquisition and maintenance of microglial identity. Transforming growth factor-beta (TGF-β) signaling is a well-established component of this developmental program, and disruption of this pathway impairs microglial maturation and homeostatic identity in both mammals and zebrafish (Butovsky et al., 2014, Arnold et al., 2019, McKinsey et al., 2025, Miserocchi et al., 2026). Radial glial progenitors have also been shown to regulate microglial TGF-β1 signaling through αVβ8-dependent local activation, implicating neural progenitor populations as an important source of instructive signals for microglial development (McKinsey et al., 2025). However, the extent to which TGF-β signaling alone accounts for the differentiation-promoting activity of the developing neural environment remains unclear. The cellular components that directly promote the transition of embryonic macrophage precursors toward a microglial state also remain incompletely defined. In particular, it is unclear whether neural progenitor cells (NPCs) directly promote this transition, how their activity compares with that of other major components of the developing brain microenvironment, such as endothelial cells (ECs), and whether macrophage precursors at different developmental stages are equivalently competent to respond to neural niche cues.

An equally important but less resolved aspect of microglial establishment is how macrophage precursors gain access to the developing neural parenchyma. In addition to parenchymal microglia, the CNS contains macrophage populations occupying its interfaces, including the meninges, perivascular spaces, and choroid plexus, which are collectively referred to as CNS-associated macrophages (CAMs) or border-associated macrophages (BAMs) (Masuda et al., 2019, Prinz et al., 2021, Dalmau Gasull et al., 2024). Microglia and non-parenchymal CNS macrophages diverge along distinct developmental trajectories, and CAM subsets subsequently acquire specialized molecular identities within defined anatomical niches (Goldmann et al., 2016, Utz et al., 2020, Masuda et al., 2022)

During embryonic development, however, macrophage phenotype, anatomical location, and eventual fate are not necessarily equivalent. CD206-positive macrophages are distributed along CNS interfaces, including the meninges and ventricular surface, and at least a subset of embryonic CD206-positive macrophages can enter the brain parenchyma and subsequently acquire microglial characteristics (Hattori et al., 2023). Thus, CD206 expression or localization at CNS interfaces during embryogenesis does not necessarily denote commitment to a mature CAM fate, and the extent to which these macrophage populations contribute to the developing microglial pool remains incompletely understood.

Several anatomical routes could potentially allow macrophage precursors to access the developing brain, including routes associated with the ventricular surface, meninges, and developing vasculature. We previously identified CD206-positive intraventricular macrophages that enter the embryonic pallium from the ventricular surface and subsequently acquire microglial characteristics, demonstrating that the embryonic ventricle can serve as a source of microglial precursors (Hattori et al., 2023). However, how the contribution of ventricular entry compares with that of other brain interfaces remains unclear. The meningeal surface represents a particularly plausible alternative route because macrophages are abundant immediately adjacent to the developing neural tissue. Recent work further suggested that early microglial progenitors can colonize the embryonic CNS from surrounding mesenchymal regions by migrating along the extracellular matrix-rich pial surface through integrin-dependent mechanisms (Petry et al., 2026). Whether meningeal entry persists beyond this early period, how it varies across anatomical regions, and what structural features of the pial boundary regulate macrophage access to the neural tissue remain unresolved. The pial basement membrane forms a specialized extracellular matrix boundary between the meninges and neural parenchyma and undergoes substantial organization during cortical development (Halfter et al., 2002, Decimo et al., 2012, Como et al., 2023), raising the possibility that its developmental maturation restricts meningeal entry.

Here, we investigated how macrophage entry into the embryonic telencephalon and their acquisition of microglial properties were developmentally regulated. We found that macrophage entry occurred through distinct routes at different embryonic stages and was associated with regional differences in pial basement membrane organization. We further found that NPCs promoted a transition toward microglial properties through close cellular proximity and, in part, TGF-β receptor signaling, and that macrophage responsiveness to these cues changed with developmental stage. Together, these findings suggest that developmentally regulated macrophage entry into the brain and stage-dependent responsiveness to local neural niche signals jointly shape microglial establishment.

## Results

### 1. Meningeal macrophage entry is restricted to an early developmental window with a regional bias

In our previous study, we showed that macrophage entry from the ventricular surface was most prominent around E12.5 and declined at later embryonic stages (Hattori et al., 2023). Because the meninges have also been proposed as a potential route of microglial colonization (Boya et al., 1991, Cuadros and Navascues, 2001, Hattori, 2023, Petry et al., 2026), we first asked whether macrophages directly enter the developing telencephalon from the meningeal surface.

To visualize macrophage behavior directly at the meningeal interface, we performed live imaging of organotypic sagittal brain slices prepared from *Cx3cr1-GFP* mice (Jung et al., 2000). To assess possible regional differences along the rostrocaudal axis, the pallium was divided into four regions, from rostral (Region 1) to caudal (Region 4) (**Fig. 1A**). CX3CR1-GFP-positive macrophages at the meningeal surface were tracked for 8 h and classified as Infiltrated, Transiently infiltrated, or Stayed (**Fig. 1B**). Cells that remained in the parenchyma for ≥4 h after entry were classified as Infiltrated, whereas those that remained for <4 h were classified as Transiently infiltrated; cells that remained within the meningeal compartment throughout the imaging period were classified as Stayed.

**Figure 1.**
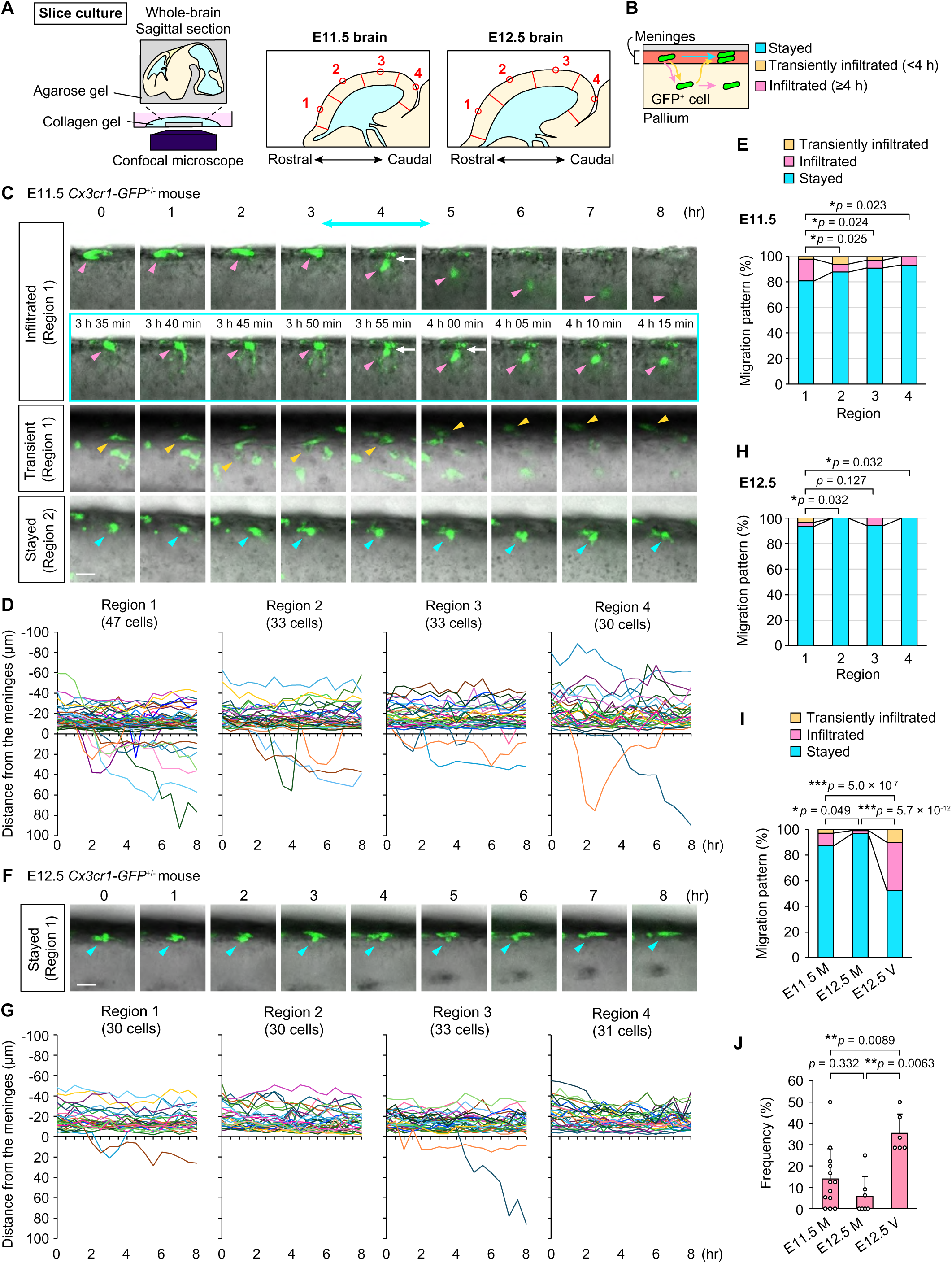
Meningeal macrophage entry is restricted to an early developmental window with a regional bias. (**A**) Schematic illustration of the organotypic slice-culture and live-imaging approach used to track macrophage migration at the meningeal surface of the embryonic pallium. (**B**) Schematic illustration of the three migration patterns used for classification: Infiltrated, Transiently infiltrated, and Stayed. (**C**) Representative time-lapse images of CX3CR1-GFP-positive macrophages at E11.5, showing cells classified as Infiltrated or Transiently infiltrated (Transient) in Region 1 and Stayed in Region 2. The cyan-boxed sequence below shows the Infiltrated cell at 5-min intervals from 3 h 35 min to 4 h 15 min. Arrowheads indicate the position of the CX3CR1-GFP-positive cell over time. White arrows indicate the cell squeezing into the brain parenchyma from the meninges. (**D**) Migration trajectories of meningeal CX3CR1-GFP-positive macrophages during the 8-h observation period, shown separately for Regions 1–4 at E11.5. (**E**) Distribution of the three migration patterns across Regions 1–4 at E11.5. Statistical significance was assessed using Pearson’s chi-squared test. (**F**) Representative time-lapse images of a CX3CR1-GFP-positive macrophage classified as Stayed in Region 1 at E12.5. (**G**) Migration trajectories of meningeal CX3CR1-GFP-positive macrophages during the 8-h observation period, shown separately for Regions 1–4 at E12.5. (**H**) Distribution of the three migration patterns across Regions 1–4 at E12.5. Statistical significance was assessed using Pearson’s chi-squared test. (**I**) Comparison of migration-pattern distributions among E11.5 meninges (E11.5 M), E12.5 meninges (E12.5 M), and the E12.5 ventricular surface (E12.5 V) after pooling data across all analyzed regions. E12.5 V data were reanalyzed from Hattori et al. (2023). Statistical significance was assessed using Pearson’s chi-squared test. (**J**) Frequency of cells classified as Infiltrated in each embryo for E11.5 M, E12.5 M, and the E12.5 V. E12.5 V data were reanalyzed from Hattori et al. (2023). Statistical significance was assessed using the two-sided Steel–Dwass test. Data are presented as the mean, with SD shown only in the positive direction. Scale bars, 20 μm (**C, F**). See also **Video S1** showing macrophage behavior across Regions 1–4 at E11.5 and E12.5.

At E11.5, macrophage entry from the meningeal surface was readily observed and showed a clear rostral bias. Infiltrated cells were most frequent in Region 1, whereas entry events were less frequent and occurred at similarly low levels across Regions 2–4 (**Fig. 1C–E**; **Video S1**). Representative time-lapse imaging of an infiltrating macrophage in Region 1 revealed a sequential entry process: the cell first extended a protrusion toward the parenchyma, followed by marked constriction of the cell body as it passed through a narrow point at the tissue boundary, and subsequently translocated into the parenchyma. Higher-temporal-resolution imaging around the entry event, acquired at 5-min intervals, further resolved these morphological changes (**Fig. 1C**, middle row). These observations indicate that meningeal macrophage entry at E11.5 is regionally biased toward the rostral pallium.

We next examined whether macrophage entry through the meningeal route persisted at E12.5. In contrast to E11.5, macrophage entry from the meningeal surface was rarely observed across the rostrocaudal axis at E12.5 (**Fig. 1F–H**; **Video S1**). The majority of tracked macrophages remained within the meningeal compartment, whereas only a small proportion showed sustained or transient parenchymal entry. Thus, macrophage entry from the meningeal surface markedly declined by E12.5 and was no longer prominent even in the rostral region.

To place the meningeal entry observed at E11.5 in the context of the ventricular route, we reanalyzed macrophage entry data from our previous study (Hattori et al., 2023). We first compared the distribution of migration patterns among E11.5 meninges, E12.5 meninges, and the E12.5 ventricular surface. The relative proportions of Infiltrated, Transiently infiltrated, and Stayed cells differed among the three conditions, with Infiltrated cells being most frequent at the E12.5 ventricular surface and least frequent at the E12.5 meningeal surface (**Fig. 1I**). We also examined the frequency of Infiltrated cells on an embryo-by-embryo basis. Consistent with the distribution analysis, the infiltrated fraction was higher at the E12.5 ventricular surface than at the E11.5 meningeal surface, whereas E12.5 meningeal entry remained rare (**Fig. 1J**). Because the ventricular and meningeal datasets were obtained using different slice orientations, differences in absolute entry frequency should be interpreted with caution. Nevertheless, the rostrocaudal axis was systematically sampled in both analyses.

Together, these findings indicate that meningeal and ventricular interfaces support macrophage entry at distinct developmental stages. Meningeal entry was most evident at E11.5 and biased toward the rostral pallium, but became rare by E12.5, when ventricular entry was prominent.

### 2. Regional differences in pial basement membrane organization are associated with meningeal macrophage entry

The regional bias of meningeal macrophage entry at E11.5 prompted us to ask whether local differences in pial border might contribute to this regional pattern of entry. Because the pial basement membrane constitutes a major structural barrier between the meninges and the neural parenchyma, we first examined its organization in sagittal sections of E11.5 *Cx3cr1-GFP*^+/−^ brains by immunohistochemistry. GFP-positive macrophages were observed entering the pallium in the rostral region (**Fig. 2A**, green box). However, in conventional sagittal sections, the laminin-positive pial basement membrane appeared broadly similar in thickness and continuity around the entry site, making it difficult to assess local structural features of the pial surface in this orientation.

**Figure 2.**
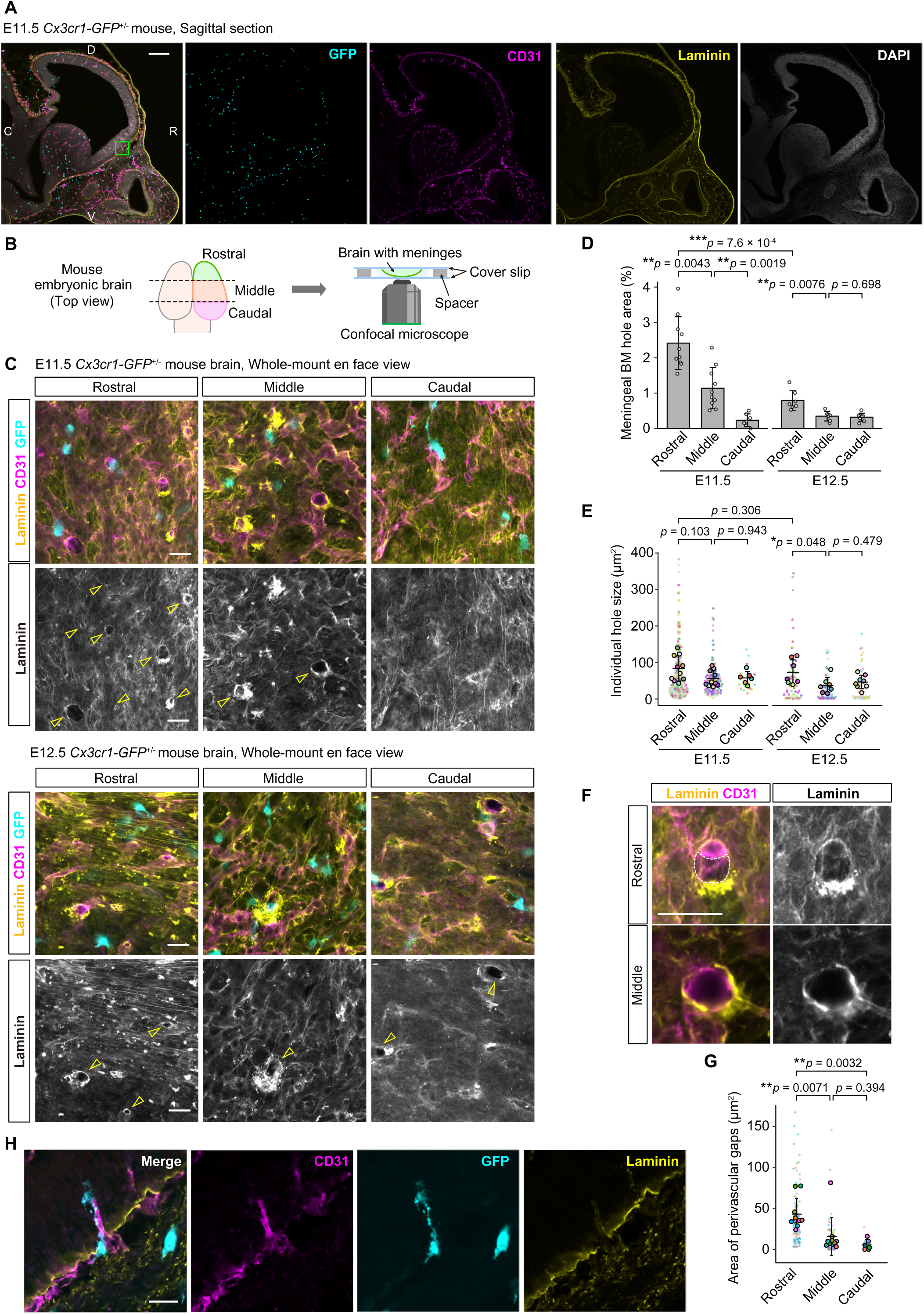
Regional differences in pial basement membrane organization are associated with meningeal macrophage entry. (**A**) Representative immunohistochemical image of a sagittal section from an E11.5 *Cx3cr1-GFP*^+/−^ brain stained for GFP, CD31, laminin, and DAPI. The green box indicates the rostral region shown at higher magnification in (**H**). D, dorsal; V, ventral; R, rostral; C, caudal. (**B**) Schematic illustration of the whole-mount immunostaining procedure for meninges attached to the brain. (**C**) Representative whole-mount immunofluorescence images of the pial surface in the rostral, middle, and caudal regions at E11.5 (top) and E12.5 (bottom), showing laminin, CD31, and CX3CR1-GFP-positive macrophages. Arrowheads indicate individual holes in the pial basement membrane. (**D**) Quantification of the frequency of pial basement membrane holes in the rostral, middle, and caudal regions at E11.5 and E12.5. Regional comparisons were performed using the two-sided Steel–Dwass test, and developmental-stage comparisons were performed using the two-sided Mann–Whitney U test (*n* = 7–10 embryos). (**E**) Quantification of the area of individual pial basement membrane holes in the rostral, middle, and caudal regions at E11.5 and E12.5. Individual holes are color-coded according to embryo, and outlined symbols indicate the mean value for each embryo. Regional comparisons were performed using the two-sided Steel–Dwass test, and developmental-stage comparisons were performed using the two-sided Mann–Whitney U test (*n* = 6–10 embryos). (**F**) Higher-magnification images of representative holes in the pial basement membrane in the rostral (top) and middle (bottom) pallium at E11.5, showing the spatial relationship of these holes to blood vessels and associated cavity-like perivascular spaces (outlined by white dashed lines). (**G**) Quantification of the area of perivascular gaps between CD31-positive vascular structures and the laminin-positive pial basement membrane in the rostral, middle, and caudal regions at E11.5. Regional comparisons were performed using the two-sided Steel–Dwass test (*n* = 6–10 embryos per region). (**H**) Higher-magnification view of the rostral region indicated by the green box in (**A**), showing a CX3CR1-GFP-positive macrophage closely associated with a pial basement membrane hole and the adjacent vascular structure at E11.5. See also **Video S2** for three-dimensional reconstruction of this region. Data are presented as the mean ± SD. Scale bars, 200 μm (**A**), 20 μm (**C, F, H**).

We therefore performed whole-mount confocal imaging to examine the pial basement membrane en face. The pial surface was immunostained for laminin to visualize the basement membrane and CD31 to label vascular endothelial structures, and basement membrane organization was compared among rostral, middle, and caudal regions at E11.5 and E12.5 (**Fig. 2B**). At E11.5, the rostral pallium contained frequent focal holes in the laminin-positive basement membrane, whereas the middle and caudal regions showed a relatively continuous laminin-positive surface. By E12.5, the pial basement membrane was largely continuous across all three regions (**Fig. 2C**).

Quantification confirmed that basement membrane holes were significantly more frequent in the rostral pallium at E11.5 than in the middle or caudal regions (**Fig. 2D**). By E12.5, hole frequency was markedly reduced overall, although the rostral region retained a tendency toward a higher frequency than the middle and caudal regions. Analysis of individual hole size showed a tendency toward larger holes in the rostral pallium, but no significant differences were detected among regions or developmental stages (**Fig. 2E**).

Closer examination of these basement membrane holes revealed that some were traversed by CD31-positive blood vessels. Higher-magnification imaging showed cavity-like spaces between the vessel surface and the surrounding laminin-positive boundary, particularly in the rostral pallium at E11.5 (**Fig. 2F**). These perivascular gaps were less evident in the middle and caudal regions and were rarely observed at E12.5, when the laminin-positive boundary more closely apposed the vessel surface. We therefore quantified the area of these gaps, excluding the area occupied by the vessels themselves. At E11.5, perivascular gap area was significantly greater in the rostral pallium than in the middle or caudal regions (**Fig. 2G**).

We next examined the spatial relationship between macrophages and these perivascular gaps using three-dimensional reconstruction of the rostral region corresponding to the area highlighted in **Fig. 2A**. Z-stack visualization showed CX3CR1-GFP-positive macrophages closely associated with these gaps, with some cells positioned between the blood vessel and the surrounding laminin-positive boundary (**Fig. 2H**; **Video S2**). Although these observations do not directly demonstrate passage of macrophages through these sites, they are consistent with the possibility that such structures provide locally permissive sites for meningeal macrophage entry.

Together, these findings show that the E11.5 rostral pallium contains more frequent basement membrane holes and larger perivascular gaps, spatially coinciding with the region in which meningeal macrophage entry was most frequently observed. The more continuous pial basement membrane and closer apposition of the basement membrane to traversing vessels at E12.5 coincide with the marked reduction in meningeal entry. These observations suggest that maturation of the pial boundary may contribute to closure of the early meningeal entry route.

### 3. NPCs promote acquisition of microglial properties in embryonic brain macrophages

We next investigated how the local neural environment promotes the acquisition of microglial properties by macrophages after their entry into the developing pallium. Although in vivo studies manipulating specific signaling pathways have provided important insights into microglial differentiation (Arnold et al., 2019, Utz et al., 2020, McKinsey et al., 2025), the direct contributions of individual cellular components of the developing neural environment remain unclear. During early brain development, CD206-positive macrophage-like cells coexist with developing microglia within the pallium. Consistent with our previous observations, CD206-positive cells accounted for approximately 53.6% of the Iba1-positive population at E12.5 and progressively declined at later stages, whereas P2RY12-positive microglia became increasingly prominent (**Fig. 3A, B; Supplemental Fig. S1**). We therefore asked whether cells within the developing neural environment directly promote this phenotypic transition.

**Figure 3.**
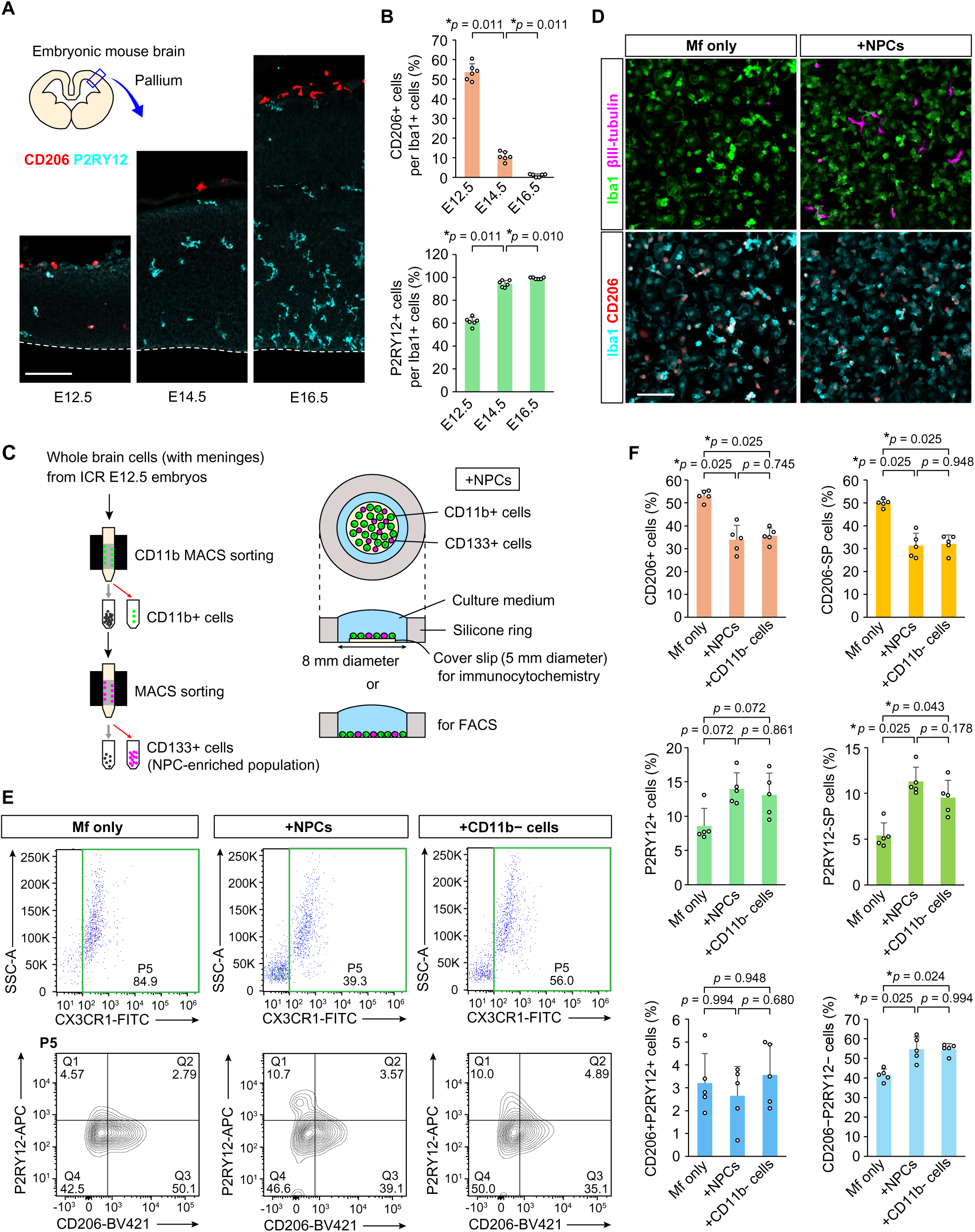
NPCs promote acquisition of microglial properties in embryonic brain macrophages. (**A**) Representative immunohistochemical images of the pallium at E12.5, E14.5 and E16.5, showing CD206 and P2RY12. See **Supplemental Fig. S1** for the corresponding Iba1 and DAPI staining. Scale bar, 100 µm. (**B**) Quantification of the proportions of CD206-positive cells (top) and P2RY12-positive cells (bottom) among Iba1-positive cells. Cell counts from multiple slides were pooled for each mouse to calculate the proportions. Each data point represents one mouse (*n* = 6 mice). Statistical significance was assessed using the two-sided Steel–Dwass test. (**C**) Schematic illustration of the isolation of CD11b-positive macrophages, CD133-positive NPC-enriched cells, and CD11b-negative cells by MACS, followed by co-culture. (**D**) Representative immunocytochemical images of macrophages cultured alone (Mf-only) or co-cultured with NPCs (+NPCs), stained for Iba1 (green) and βIII-tubulin (magenta) in the upper panels and Iba1 (cyan) and CD206 (red) in the lower panels. No detectable βIII-tubulin-positive neural-lineage cells were observed in the Mf-only condition. Scale bar, 100 µm. (**E**) Representative flow cytometric analysis of macrophages cultured alone or co-cultured with NPCs or CD11b-negative cells. P2RY12 and CD206 expression was analyzed within the CX3CR1-positive population (P5, shown in green). See also **Supplemental Fig. S2**. (**F**) Quantification of the percentages of CD206-positive, CD206-single-positive (SP), P2RY12-positive, P2RY12-SP, CD206/P2RY12-double-positive, and CD206/P2RY12-double-negative cells among total CX3CR1-positive cells (P5). Statistical significance was assessed using the Steel–Dwass test (*n* = 5 independent experiments). Data are presented as mean, with SD shown only in the positive direction.

To address this question, we established an in vitro co-culture system using embryonic macrophage populations and non-hematopoietic cells isolated from the developing brain (**Fig. 3C**). Whole brains, including the meninges, were collected from E12.5 embryos, and CD11b-positive cells were isolated by magnetic-activated cell sorting (MACS). Because the meninges cannot be reliably separated from the brain at this developmental stage, the E12.5 CD11b-positive fraction contained macrophages from multiple CNS compartments, including the meninges and ventricles, together with cells already present within the parenchyma. From the same brains, CD133-positive cells were isolated as an NPC-enriched population. CD11b-positive cells were cultured alone (Mf-only) or co-cultured with NPCs (+NPCs) for 2 days. Immunocytochemical analysis detected no βIII-tubulin-positive neural-lineage cells in the Mf-only condition, whereas such cells were readily observed in the +NPC condition, confirming minimal neural-lineage contamination in the Mf-only cultures and the successful establishment of the NPC co-culture system (**Fig. 3D**).

First, we examined whether exposure to NPCs altered macrophage phenotype by flow cytometry, using CD206 and P2RY12 to distinguish macrophage-like and microglial properties within the 7-AAD-negative, CX3CR1-positive population (P5). Co-culture with NPCs significantly reduced both the CD206-positive and CD206-single-positive populations, while significantly increasing the P2RY12-single-positive population (**Fig. 3E, F**; **Supplemental Fig. S2**). The overall proportion of P2RY12-positive cells showed a modest increase but did not reach statistical significance. The double-negative population also changed significantly, whereas the double-positive population was not significantly altered. Thus, NPC co-culture induced a clear shift away from a CD206-positive macrophage phenotype, accompanied by an increase in the P2RY12-single-positive population. However, the overall increase in P2RY12-positive cells was modest, suggesting that NPC exposure promoted a phenotypic transition rather than complete acquisition of microglial identity.

We next asked whether other non-hematopoietic brain-cell populations enhanced this phenotypic transition beyond the effect of NPCs alone. Macrophages were cultured alone, with purified NPCs, or with unfractionated CD11b-negative brain cells and analyzed under the same conditions. The CD11b-negative cell condition produced a pattern of phenotypic changes similar to that observed with purified NPCs, including significant reductions in the CD206-positive and CD206-single-positive populations and significant increases in the P2RY12-single-positive and double-negative populations (**Fig. 3E, F**; **Supplemental Fig. S2**). No significant differences were detected between the +NPC and +CD11b-negative cell conditions.

Thus, purified NPCs were sufficient to reproduce the phenotypic changes induced by the broader CD11b-negative brain-cell population, with no detectable additional effect of other non-hematopoietic cells under these culture conditions. Together, these findings identify NPCs as a major cellular component of the embryonic neural environment capable of driving embryonic macrophages away from a CD206-high phenotype.

### 4. NPC-induced macrophage phenotypic transition requires close cellular proximity and partly depends on TGF-β signaling

To further define the cellular basis of the NPC-induced phenotypic transition observed above, we compared the effects of major non-hematopoietic cell populations within the developing brain. CD133-positive and CD31-positive cells were isolated from E12.5 brains as NPC- and EC-enriched populations, respectively. CD11b-positive macrophages were cultured alone (Mf-only), with NPCs (+NPCs), with ECs (+ECs), or with both NPCs and ECs (+NPCs/ECs) for 2 days (**Fig. 4A**).

**Figure 4.**
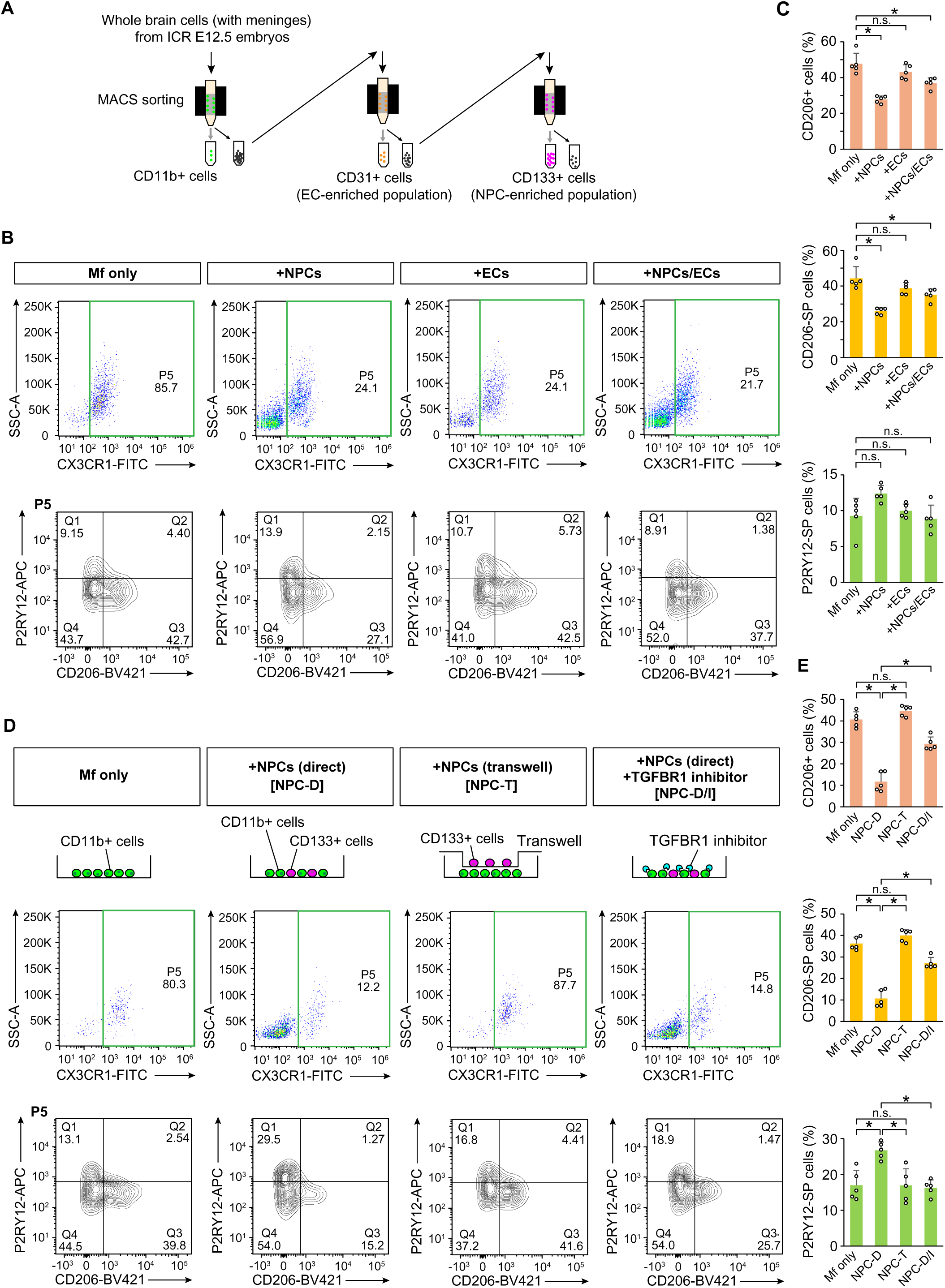
NPC-induced macrophage phenotypic transition requires close cellular proximity and partly depends on TGF-β signaling. (**A**) Schematic illustration of the sequential isolation of CD11b-positive macrophages, CD31-positive EC-enriched cells, and CD133-positive NPC-enriched cells by MACS, followed by co-culture. (**B**) Representative flow cytometric analysis of macrophages cultured alone (Mf-only), with NPCs (+NPCs), with ECs (+ECs), or with both NPCs and ECs (+NPCs/ECs). CD206 and P2RY12 expression was analyzed within the CX3CR1-positive population (P5, shown in green). See also **Supplemental Fig. S3, S4A**. (**C**) Quantification of the percentages of CD206-positive, CD206-single-positive (SP), and P2RY12-SP cells among total CX3CR1-positive cells (P5). Statistical significance was assessed using the two-sided Steel–Dwass test (*n* = 5 independent experiments). For comparisons of Mf-only with +NPCs, +ECs, and +NPCs/ECs, respectively, *p* values were 0.045, 0.784, and 0.045 for CD206-positive cells; 0.045, 0.398, and 0.045 for CD206-SP cells; and 0.075, 1.000, and 0.884 for P2RY12-SP cells. (**D**) Representative flow cytometric analysis of macrophages cultured alone (Mf-only), directly co-cultured with NPCs (NPC-D), indirectly co-cultured with NPCs using a Transwell system (NPC-T), or directly co-cultured with NPCs in the presence of the TGF-β receptor signaling inhibitor SB431542 (NPC-D/I). See also **Supplemental Fig. S4B, S5**. (**E**) Quantification of the percentages of CD206-positive, CD206-SP, and P2RY12-SP cells among total CX3CR1-positive cells (P5, shown in green). Statistical significance was assessed using the two-sided Steel–Dwass test (*n* = 5 independent experiments). For Mf-only vs. NPC-D, Mf-only vs. NPC-T, NPC-D vs. NPC-T, and NPC-D vs. NPC-D/I, respectively, *p* values were 0.045, 0.194, 0.045, and 0.045 for CD206-positive cells; 0.044, 0.285, 0.044, and 0.044 for CD206-single-positive cells; and 0.045, 1.000, 0.045, and 0.045 for P2RY12-single-positive cells. Data are presented as the mean, with SD shown only in the positive direction.

As observed in the preceding experiments, co-culture with NPCs significantly reduced both the CD206-positive and CD206-single-positive populations relative to the Mf-only condition (**Fig. 4B, C**; **Supplemental Fig. S3, S4A**). P2RY12-positive populations showed a modest tendency to increase, but these changes did not reach statistical significance. In contrast, co-culture with ECs did not significantly affect the populations examined relative to the Mf-only condition.

We also examined simultaneous co-culture with NPCs and ECs. To maintain the same total number of added cells across conditions, the numbers of NPCs and ECs in the +NPCs/ECs condition were each reduced to half of those used in the corresponding single-cell-type conditions. The +NPCs/ECs condition produced a pattern of phenotypic changes similar to that observed with NPCs alone, including significant reductions in the CD206-positive and CD206-single-positive populations, without significant changes in P2RY12-positive populations. Thus, under these experimental conditions, ECs did not produce a detectable phenotypic effect on their own and did not provide an additional effect beyond that induced by NPCs. Together, these findings indicate that NPCs, but not ECs, are a major non-hematopoietic cell population capable of driving embryonic macrophages away from a CD206-high phenotype.

We next asked whether the NPC-induced phenotypic transition required close cellular proximity. Previous studies have implicated αVβ8-dependent local activation of TGF-β by neural progenitor populations in microglial development, raising the possibility that short-range interactions within the neural niche contribute to the effects observed here (Arnold and Betsholtz, 2013, McKinsey et al., 2025). CD11b-positive macrophages and CD133-positive NPCs were therefore isolated from E12.5 brains and cultured under four conditions: macrophages alone (Mf-only), direct co-culture with NPCs (Direct), indirect co-culture with NPCs physically separated by a permeable Transwell membrane (Transwell), or direct co-culture with NPCs in the presence of the TGF-β receptor signaling inhibitor SB431542 (Direct + SB431542). After 2 days of culture, CD206 and P2RY12 expression within the CX3CR1-positive population was assessed by flow cytometry.

Direct co-culture with NPCs significantly reduced both the CD206-positive and CD206-single-positive populations and significantly increased the P2RY12-positive and P2RY12-single-positive populations relative to the Mf-only condition (**Fig. 4D, E**; **Supplemental Fig. S4B, S5**). In contrast, when macrophages and NPCs were physically separated in the Transwell condition, these changes were largely lost, and the distribution of CD206- and P2RY12-expressing populations remained similar to that observed in the Mf-only condition. These findings indicate that close cellular proximity is required for the full NPC-induced phenotypic transition.

Inhibition of TGF-β receptor signaling during direct co-culture partially attenuated the NPC-induced changes. The reduction in CD206-positive populations was less pronounced in the presence of SB431542, and the increase in P2RY12-positive populations was also partially suppressed (**Fig. 4D, E**; **Supplemental Fig. S4B, S5**). Thus, TGF-β receptor signaling contributes to, but does not fully account for, the phenotypic changes induced by NPCs.

Together, these findings indicate that the NPC-induced transition of embryonic macrophages away from a CD206-high state requires close cellular proximity and is mediated, at least in part, by TGF-β receptor signaling. The incomplete suppression by SB431542 further suggests that additional short-range signals within the neural niche contribute to this process.

### 5. Macrophage competence to acquire microglial properties changes with developmental stage

Because embryonic CNS macrophages undergo substantial developmental changes over time (Matcovitch-Natan et al., 2016, Li et al., 2019, Utz et al., 2020), we next asked whether their responsiveness to NPC-derived cues changes with developmental stage. To address this question, CD11b-positive macrophages were isolated from the meninges at E13.5 or E17.5 and co-cultured under identical conditions with the same E12.5 CD133-positive NPC population. E13.5 was selected as the earliest stage at which the meninges could be reproducibly dissected, allowing macrophages from the same anatomical compartment to be compared across developmental stages. E13.5 meningeal macrophages showed a pronounced phenotypic response to NPC co-culture. In the Mf-only condition, most cells were CD206 positive, whereas only a small proportion expressed P2RY12 (**Fig. 5A, B**; **Supplemental Fig. S6**). Following NPC co-culture, the CD206-positive population decreased from 88.8% to 50.4%, whereas the P2RY12-positive population increased from 9.0% to 23.2%. Notably, the P2RY12-positive/CD206-negative single-positive population increased to 19.8%, indicating robust acquisition of microglial properties.

**Figure 5.**
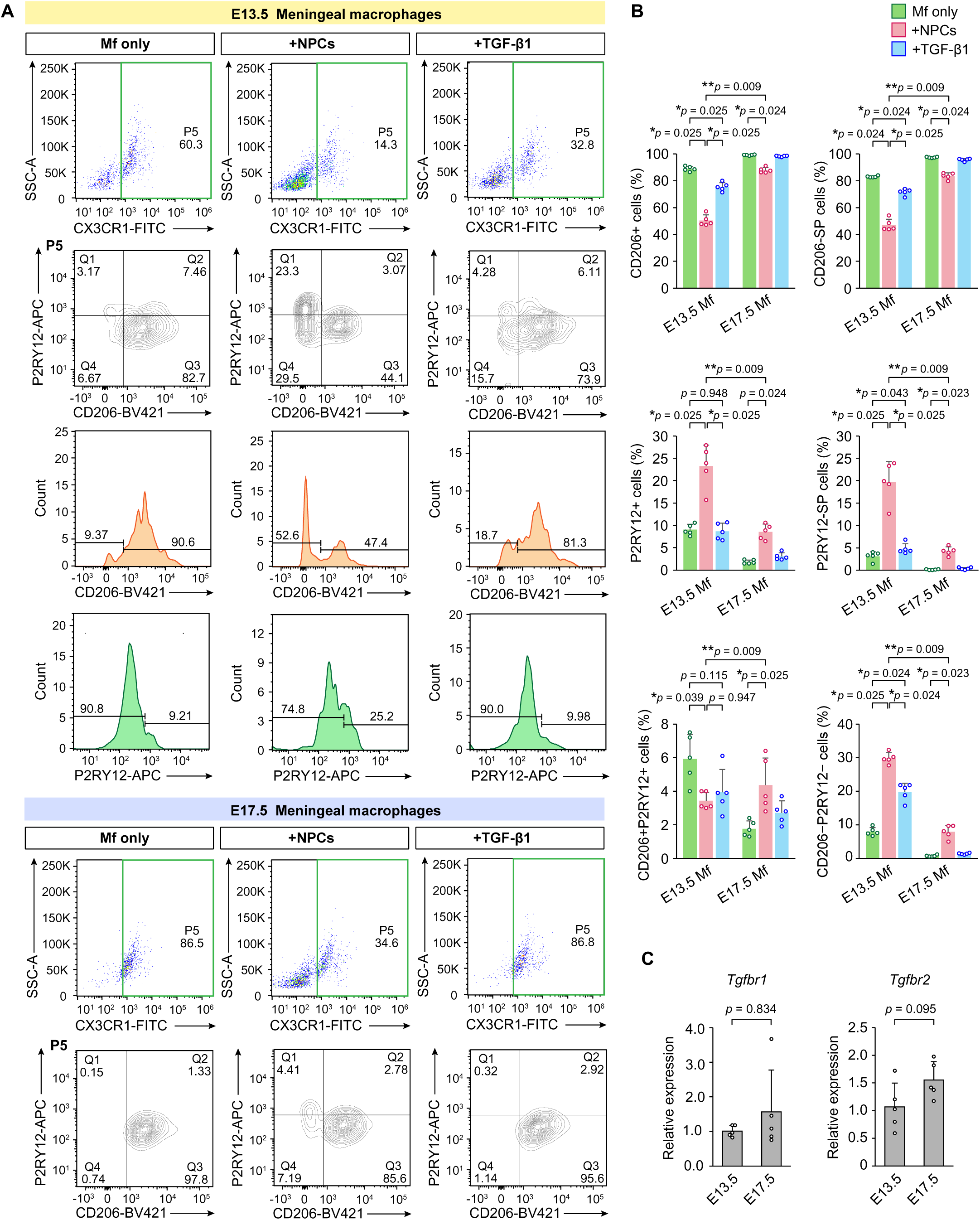
Macrophage competence to acquire microglial properties changes with developmental stage. (**A**) Representative flow cytometric analysis of meningeal macrophages isolated from E13.5 or E17.5 embryos and cultured alone (Mf-only), co-cultured with an E12.5 CD133-positive NPC-enriched population (+NPCs), or cultured with TGF-β1 (+TGF-β1). P2RY12 and CD206 expression was analyzed within the CX3CR1-positive population (P5, shown in green). See also **Supplemental Fig. S6, S7**. (**B**) Quantification of the percentages of CD206-positive, CD206-single-positive, P2RY12-positive, P2RY12-single-positive, and CD206/P2RY12-double-positive cells among total CX3CR1-positive cells (P5, shown in green). Statistical significance was assessed using the two-sided Steel–Dwass test unless otherwise indicated (*n* = 5 independent experiments). (**C**) Quantitative PCR analysis of *Tgfbr1* and *Tgfbr2* expression in E13.5- and E17.5-derived meningeal macrophages. Expression levels were normalized to *Hprt*. Data are presented as the mean, with SD shown only in the positive direction.

In contrast, E17.5 meningeal macrophages responded much less strongly to the same NPC population. CD206-positive cells remained predominant following NPC co-culture (99.2% in Mf-only versus 87.9% with NPCs), and the increase in P2RY12-positive cells was modest (1.9% to 8.6%) (**Fig. 5A, B**; **Supplemental Fig. S7**). Importantly, the P2RY12-positive/CD206-negative single-positive population reached only 4.2%, compared with 19.8% in E13.5 macrophages. Because E13.5 and E17.5 macrophages were exposed to the same NPC population under otherwise identical culture conditions, these findings indicate that the capacity of meningeal macrophages to acquire microglial properties in response to neural niche cues changes substantially with developmental stage.

We next asked whether recombinant TGF-β1 alone could reproduce the phenotypic changes induced by NPCs. E13.5 and E17.5 meningeal macrophages were therefore analyzed in parallel under Mf-only, +NPC, and +TGF-β1 conditions. Recombinant TGF-β1 had only a limited effect on P2RY12 expression at either developmental stage. In E13.5 macrophages, the P2RY12-positive population remained close to the Mf-only level following TGF-β1 treatment (8.7%), in contrast to the increase to 23.2% induced by NPC co-culture. Similarly, in E17.5 macrophages, P2RY12-positive cells increased only slightly following TGF-β1 treatment (1.9% to 3.1%), compared with 8.6% following NPC co-culture (**Fig. 5A, B**; **Supplemental Figs. S6 and S7**). Thus, recombinant TGF-β1 alone was insufficient to reproduce the NPC-induced increase in P2RY12 expression under these culture conditions.

To determine whether the developmental difference in NPC responsiveness could be explained by altered expression of TGF-β receptors, we examined *Tgfbr1* and *Tgfbr2* expression in E13.5 and E17.5 meningeal macrophages. Neither *Tgfbr1* nor *Tgfbr2* expression differed significantly between the two developmental stages (**Fig. 5C**). These results suggest that differences in *Tgfbr1* and *Tgfbr2* mRNA expression are unlikely to fully account for the stage-dependent difference in responsiveness to NPC-derived cues.

Together, these findings indicate that meningeal macrophages differ in their competence to acquire microglial properties depending on developmental stage, with E13.5 macrophages showing a stronger response to NPC co-culture than E17.5 macrophages. Recombinant TGF-β1 alone was insufficient to reproduce this response, suggesting that additional NPC-associated cues contribute to the phenotypic transition.

## Discussion

In this study, we identified two developmentally regulated processes that together shape microglial establishment in the embryonic brain: the ability of macrophage precursors to respond to neural niche signals and their anatomical access to the developing telencephalon. NPCs promoted a phenotypic transition of embryonic macrophages, with earlier-stage meningeal macrophages showing a particularly robust acquisition of microglial properties, whereas later-stage macrophages were less responsive to the same NPC population. In parallel, macrophage entry occurred through temporally distinct ventricular and meningeal routes. These findings suggest that microglial establishment depends on macrophage developmental state, local neural niche signals, and the developmental regulation of macrophage entry into the telencephalon.

Neural progenitor populations have previously been implicated in microglial positioning through CXCL12 and in microglial maturation through local regulation of TGF-β signaling (Arno et al., 2014, Hattori et al., 2020, McKinsey et al., 2025). Our findings extend these observations by showing that the macrophage phenotypic transition induced by NPCs depended strongly on close cellular proximity. Pharmacological inhibition of TGF-β receptor signaling attenuated this effect, indicating that TGF-β signaling forms part of the local NPC–macrophage interaction. However, exogenous active TGF-β1 alone produced only a modest response and failed to reproduce the effect of NPC co-culture. Thus, local activation of TGF-β is unlikely to fully account for the effect of NPCs. Instead, TGF-β may function as one component of a spatially organized combination of growth factors, membrane-associated ligands, extracellular matrix components, or other short-range cues provided by NPCs.

A particularly important implication of our findings is that the extent to which macrophages acquire microglial properties in an NPC-containing environment may depend on their developmental state. E13.5 and E17.5 meningeal macrophages were exposed to the same E12.5 NPC population, yet earlier-stage macrophages underwent a substantially stronger transition toward a P2RY12-positive/CD206-negative phenotype. *Tgfbr1* and *Tgfbr2* transcript levels did not differ detectably between the two developmental stages, suggesting that differences in receptor gene expression alone do not explain the distinct phenotypic responses. Developmental changes in intracellular signaling or transcriptional competence may instead contribute to this difference. TGF-β signaling converges on SMAD-dependent transcriptional programs that are required for acquisition of microglial identity, including transcriptional programs involving SALL1 (Brioschi et al., 2023). In addition, microglial maturation is accompanied by progressive remodeling of the enhancer landscape; IRF8, together with SALL1 and PU.1, promotes stepwise increases in chromatin accessibility that precede establishment of mature microglial transcriptional programs (Saeki et al., 2024). Although these mechanisms have not been examined directly in our embryonic macrophage populations, they raise the possibility that stage-dependent differences in the macrophage phenotypic response to NPCs reflect altered intracellular signaling or epigenetic competence rather than differences in receptor expression alone. These cell-intrinsic developmental programs are likely to interact with local tissue context. Embryonic microglia also adopt spatially restricted cellular states associated with specific anatomical and morphogenetic environments (Lawrence et al., 2024), further supporting the idea that developmental state and tissue context jointly shape microglial identity.

Developmental regulation was also evident at the level of anatomical access. We previously showed that CD206-positive macrophages within the ventricular lumen can enter the developing pallium and subsequently acquire microglial characteristics (Hattori et al., 2023, Hattori, 2025). In the present study, meningeal entry was detected in the rostral pallium at E11.5 and became rare by E12.5. Reanalysis of our previously acquired imaging data showed prominent ventricular entry at E12.5. These observations identify an early, transient meningeal entry route alongside the ventricular route observed at E12.5. Recent work showing that early microglial progenitors migrate along the extracellular matrix-rich pial surface supports an important role for the meningeal interface during initial brain colonization (Petry et al., 2026). Our findings added a temporal and regional dimension to this model by indicating that macrophage entry across the pial interface was spatially and temporally restricted, representing a transient developmental opportunity that narrowed rapidly as the developing pallium matured.

The spatial correspondence between meningeal entry and pial basement membrane organization provided a potential structural basis for this restricted pattern of meningeal entry. At E11.5, cell-sized discontinuities were selectively enriched in the rostral pial basement membrane, where meningeal entry was observed, whereas the boundary was more continuous in middle and caudal regions and became largely continuous by E12.5. Whole-mount three-dimensional imaging further revealed macrophages positioned in close association with these local openings. Although these observations do not directly demonstrate passage of macrophages through the discontinuities, they suggest that such sites may provide locally permissive regions for entry.

Basement membranes are increasingly recognized as spatially heterogeneous barriers that can contain preferential sites for leukocyte passage (Nourshargh et al., 2010, Li, 2025). In venular basement membranes, matrix protein low-expression regions serve as preferred exit sites for neutrophils (Wang et al., 2006), and subsequent studies demonstrated that such regions are present across multiple tissues and function as permissive “gates” for leukocyte transmigration (Voisin et al., 2010). Macrophages can likewise traverse native basement membranes by exploiting pre-existing pores, including through protease-independent, actomyosin-dependent mechanisms (Bahr et al., 2022). By contrast, endothelial tip cells actively remodel the pial basement membrane through matrix metalloproteinase-25 (MMP25)-dependent proteolysis during initial brain vascularization (Schevenels et al., 2024). Together, these studies suggest that basement membranes can regulate cell entry not simply as uniform physical barriers, but through local differences in structural permissiveness. Our findings raise the possibility that embryonic macrophages similarly take advantage of transient pre-existing discontinuities in the developing pial basement membrane.

The capacity of embryonic interface-associated macrophages to acquire microglial properties has implications for how their identity is interpreted. Transplantation studies have shown that both cellular origin and the CNS environment contribute to the acquisition of microglial identity (Bennett et al., 2018, Ayata et al., 2018). During embryogenesis, residence outside the neural parenchyma may therefore not imply irreversible commitment to a CAM identity. We previously showed that CD206-positive intraventricular macrophages can enter the pallium and subsequently acquire microglial properties (Hattori et al., 2023). Our findings also highlight a distinction between the anatomical identity of embryonic CNS macrophages and their developmental potential. The acquisition of microglial properties by meningeal macrophages in NPC co-culture suggests that residence outside the neural parenchyma does not necessarily limit their ability to respond to neural niche signals. Anatomical position and marker expression may therefore describe their current phenotype without fully defining their capacity to acquire microglial properties.

Although our findings link developmental differences in macrophage phenotypic transition with anatomical opportunities for brain entry, the mechanisms underlying these processes and their relationship in vivo remain unresolved. The molecular combination through which NPCs promote macrophage phenotypic transition remains undefined, and the limited effect of recombinant TGF-β1 suggests that additional short-range signals will need to be identified. Likewise, the molecular basis of the developmental change in macrophage competence remains unknown and will require direct comparison of intracellular signaling, transcriptional programs, and chromatin states between early- and late-stage macrophages. Although the spatial correspondence between pial basement membrane discontinuities and meningeal entry was striking, causal manipulation of basement membrane organization will be required to determine whether maturation of this boundary directly restricts macrophage access. Finally, lineage-resolved approaches will be necessary to quantify the relative contributions of ventricular, meningeal, and other potential entry routes to the mature microglial population.

In summary, our findings suggest that microglial establishment requires the coincidence of two permissive conditions: a macrophage state competent to respond to neural niche signals and a tissue state permissive for entry into the neural parenchyma. Developmental changes in either condition could restrict when and where embryonic macrophages contribute to the developing microglial pool. Thus, microglial establishment is restricted to developmental periods when macrophages are competent to respond to neural cues and brain interfaces permit their entry into the developing telencephalon.

## Acknowledgments

We thank Makoto Masaoka, Ikuko Mizuno, and Namiko Noguchi (Department of Anatomy and Cell Biology, Nagoya University Graduate School of Medicine) for their technical assistance. We wish to acknowledge the Division for Medical Research Engineering, Nagoya University Graduate School of Medicine, for technical support. This work was supported by Grants-in-Aid for Transformative Research Areas (A) (JP21H05624 [Y.H.]; JP23H04161 [Y.H.]), Grant-in-Aid for Scientific Research (B) (JP23K27349 [Y.H.]; JP26K02191 [Y.H.]; JP24K02205 [T.M.]), FOREST program (JPMJFR214C [Y.H.]), Grant-in-Aid for Research Activity Start-up (JP25K23790 [T.S.]), grants from AMED-ASPIRE program (JP23jf0126004 [Y.H.]). It was also supported by grants from the Mochida Memorial Foundation for Medical and Pharmaceutical Research, the Chugai Foundation for Innovative Drug Discovery Science, and the Takeda Science Foundation [Y.H.].

## Author contributions

Conceptualization: Y.H.; Methodology: M.O., T.S., Z.Y., T.M., and Y.H.; Investigation: M.O., T.S., Z.Y., and Y.H.; Writing – Original Draft, Y.H.; Writing – Review & Editing, M.O., T.S., T.M., and Y.H.; Visualization: Y.H.; Project administration: Y.H.; Funding acquisition: T.S., T.M., and Y.H. All authors approved the final version of the manuscript and agree to be accountable for all aspects of the work.

## Declaration of interests

The authors declare no competing interests.

## Materials and Methods

### Resource availability

#### Lead contact

All unique/stable reagents generated in this study are available from the Lead Contact without restriction. Further information or request for resources and reagents should be directed to and will be fulfilled by Yuki Hattori.

#### Materials availability

This study did not generate new unique reagents.

#### Data and code availability

The source data are provided as a Source Data file. The data have also been deposited in Mendeley Data (doi: 10.17632/n33kdpgcbp.1) and will be publicly available upon publication. For all other inquiries, please contact the corresponding author.

### Experimental model and subject details

ICR mice were purchased from Japan SLC (Shizuoka, Japan). *Cx3cr1-GFP* mice (Stock No. 005582, RRID: IMSR_JAX:005582) were purchased from Jackson Laboratory (Jung et al., 2000). All mice were housed under specific-pathogen-free conditions at Nagoya University. The animal experiments were conducted according to the Japanese Act on Welfare and Management of Animals, Guidelines for Proper Conduct of Animal Experiments (published by the Science Council of Japan), Fundamental Guidelines for Proper Conduct of Animal Experiment and Related Activities in Academic Research Institutions (published by Ministry of Education, Culture, Sports, Science and Technology, Japan). All protocols for animal experiments were approved by the Institutional Animal Care and Use Committee of Nagoya University (No. 29006).

To obtain *Cx3cr1-GFP*^+/-^ embryos (heterozygous), male *Cx3cr1-GFP* homozygous mice were mated with female ICR mice. The day when the vaginal plug was detected was counted as E0.5. Both male and female embryos were used, and similar results were obtained.

### Method details

#### Live imaging in cultured cortical slices

To obtain cortical slices with intact meninges, whole forebrains isolated from E11.5 to E17.5 male and female *Cx3cr1-GFP*^+/-^ mice were embedded in 2% agarose gel and sliced coronally or sagittally (400 µm thickness) using a vibratome (**Fig. 1**). The agarose-embedded slices were placed on 3.5 cm glass-bottom dishes (27 mm glass diameter; Iwaki [AGC Techno Glass Co., Ltd.], Cat# 3910-035) and then mounted in collagen gel (Cellmatrix Type I-A [Collagen, Type I, 3 mg/mL, pH 3.0]; Nitta Gelatin, Cat#KP-2100) (Hattori et al., 2020). After the collagen gel solidified, 2–3 ml of the culture medium was added. The composition of the culture medium was as follows: D-MEM/F12 (Sigma–Aldrich, Cat# D2906) supplemented with 5% fetal bovine serum (FBS; Invitrogen), 5% horse serum (HS; Invitrogen), N-2 Supplement (Thermo Fisher Scientific, Cat# 17502001), B-27 Supplement without vitamin A (Thermo Fisher Scientific, Cat# 12587010), and penicillin/streptomycin (50 U/ml, each; Meiji Seika Pharma Co., Ltd.).

For cell tracking analysis, time-lapse imaging was performed using a CV1000 confocal microscope (Yokogawa). The on-stage culture chamber was filled with 40% O and 5% CO . CX3CR1 macrophages initially located in the meninges were tracked at 5-minute intervals for 8 hours (*n* = 13 embryos at E11.5 and 7 embryos at E12.5). To determine the frequency of macrophage entry from the lateral ventricle at E12.5, we reanalyzed our previously published time-lapse imaging dataset (*n* = 6 embryos) (Hattori et al., 2023).

Migration patterns were classified into three groups based on cell behavior during the 8-h observation period. Cells that entered the pallial parenchyma and remained there for ≥4 h were classified as Infiltrated, regardless of whether they subsequently returned to the meningeal surface. Cells that entered the pallial parenchyma but returned to the meningeal surface within <4 h were classified as Transiently infiltrated, whereas cells that remained in the meninges throughout the observation period were classified as Stayed. For imaging of sagittal sections, the pallium was divided into four equal regions along the rostrocaudal axis, and the proportions of cells exhibiting each migration pattern were quantified in each region.

#### Cell isolation and cultures

For experiments shown in **Fig. 3 and 4**, whole brains, including the meninges, were dissected from E12.5 embryos, mechanically minced, and incubated in 0.05% trypsin–EDTA at 37°C for 5 min. The tissues were dissociated by pipetting and passed through a 40-μm cell strainer to obtain single-cell suspensions. Cells were sequentially separated into CD11b cells, CD31 endothelial cells (ECs), and CD133 neural progenitor cells (NPCs) by magnetic-activated cell sorting (MACS), using PBS containing 0.5% FBS as the MACS buffer. First, the entire cell suspension was labeled with CD11b (Microglia) MicroBeads (human and mouse; Miltenyi Biotec, Cat# 130-093-634), and CD11b cells were enriched by two successive rounds of positive selection using MACS columns. The resulting CD11b fraction was then labeled with CD31 MicroBeads (mouse; Miltenyi Biotec, Cat# 130-097-418), and CD31 ECs were isolated by one round of positive selection. Finally, the CD11b CD31 fraction was labeled with Anti-Prominin-1 (CD133) MicroBeads (mouse; Miltenyi Biotec, Cat# 130-092-333), and CD133 NPC-enriched cells were isolated by one round of positive selection.

For Transwell co-culture experiments, E12.5 CD11b macrophages (5 × 10 cells/well) were seeded in the lower chamber, and E12.5 CD133 NPC-enriched cells (5 × 10 cells/insert) were seeded in Transwell inserts with a porous membrane that permitted the exchange of soluble factors while preventing direct cell–cell contact (Millicell, Millipore, 0.4-μm pore size, 30-mm diameter, Cat# PICM0RG50). Macrophages and NPCs were cultured for 2 days under otherwise identical conditions to those used for direct co-culture. After culture, cells from the lower chamber were collected and analyzed by flow cytometry for CD206 and P2RY12 expression within the CX3CR1-positive population.

For the developmental-stage comparison shown in **Fig. 5**, meninges were dissected from E13.5 or E17.5 brains and dissociated as described above. Meningeal cell suspensions were labeled with CD11b (Microglia) MicroBeads, and CD11b macrophages were isolated by one round of positive selection. For co-culture experiments, these E13.5 or E17.5 meningeal macrophages were cultured with CD133 NPC-enriched cells isolated from E12.5 brains as described above.

Each isolated cell population was subsequently resuspended in DMEM/F12. Cells were seeded within silicone rings (8-mm inner diameter) placed on polyethyleneimine (PEI)-coated culture dishes. The following three culture conditions were established: CD11b cells alone (1 × 10 cells/well), coculture with NPCs (5 × 10 CD11b cells and 5 × 10 CD133 cells/well), and coculture with ECs (5 × 10 CD11b cells and 5 × 10 CD31 cells/well).

For the developmental-stage comparison shown in **Fig. 5**, fewer meningeal macrophages were available; therefore, cultures were established at 70% of the total cell number used in **Fig. 3 and 4**. CD11b meningeal macrophages were cultured alone (7.0 × 10 cells/well) or co-cultured with E12.5 CD133 NPCs (3.5 × 10 CD11b cells and 3.5 × 10 CD133 cells/well), maintaining the same 1:1 macrophage-to-NPC ratio. ECs were not included in these experiments.

All cells were cultured for 2 days at 37°C in a humidified atmosphere containing 5% CO in DMEM/F12 (Sigma–Aldrich, Cat# D2906) supplemented with 10% FBS (Invitrogen), N-2 Supplement (Thermo Fisher Scientific, Cat# 17502001), B-27 Supplement without vitamin A (Thermo Fisher Scientific, Cat# 12587010), and penicillin–streptomycin (50 U/mL each; Meiji Seika Pharma Co., Ltd.).

#### Immunocytochemistry

Cells were cultured on 5-mm-diameter circular coverslips precoated with polyethyleneimine (PEI), which were placed inside 8-mm-diameter silicone rubber wells. After culture, cells were fixed with 4% paraformaldehyde (PFA) for 10 min at room temperature and incubated overnight at 4°C with the following primary antibodies: goat anti-CD206 polyclonal antibody (1:300; R&D Systems, Cat# AF2535, RRID:AB_2063012), chicken anti-Iba1 antibody (1:1,000; Synaptic Systems, Cat# 234 009, RRID:AB_2891282), and rabbit anti-βIII-tubulin (Tuj1) polyclonal antibody (1:1,000; BioLegend, Cat# MRB-435P, RRID:AB_663339) (**Fig. 3D**).

After washing, cells were incubated with appropriate Alexa Fluor 488-, Alexa Fluor 546-, or Alexa Fluor 647-conjugated secondary antibodies (1:1,000; Invitrogen, Cat# A78948, RRID:AB_2921070; Cat# A11056, RRID:AB_2534103; Cat# A31573, RRID:AB_2536183), together with DAPI (1:1,000; Sigma–Aldrich, Cat# D9542). After staining, cells were mounted in mounting medium and imaged using an AX confocal microscope (Nikon).

#### Immunohistochemistry

Brains were fixed in 4% PFA, cryoprotected in 20% sucrose, and then sectioned (16 µm) on a cryostat. Sections were incubated with the following primary antibodies overnight at 4°C: goat anti-CD206 polyclonal antibody (1:300, R&D Systems, Cat# AF2535, RRID:AB_2063012), goat anti-CD31 polyclonal antibody (1:300, R&D Systems, Cat# AF3628, RRID:AB_2161028), chicken anti-GFP polyclonal antibody (1:1000; Aves Labs, Cat# GFP-1020, RRID:AB_10000240), chicken anti-Iba1 monoclonal antibody (1:1,000, Synaptic Systems, Cat# 234 009, RRID:AB_2891282), guinea pig anti-Laminin polyclonal antibody (1:1,000, Wako, Cat# 010-28821, RRID:AB_3754827), and rabbit anti-P2RY12 polyclonal antibody (1:500, AnaSpec, Cat# 55043A, RRID: AB_2298886).

After washing, the sections were incubated with Alexa Fluor 488-, Alexa Fluor 546-, or Alexa Fluor 647-conjugated secondary antibodies (1:1,000; Invitrogen, Cat# A11055, RRID:AB_2534102; Cat# A11056, RRID:AB_2534103; Cat# A78948, RRID:AB_2921070) or Alexa Fluor 647-conjugated anti-guinea pig IgG antibody (1:1,000; Jackson ImmunoResearch Laboratories, Cat# 706-606-148, RRID:AB_2340477), together with DAPI (1:1,000; Sigma–Aldrich, Cat# D9542). For P2RY12 staining, antigen retrieval was performed using HistoVT (Nacalai Tesque) at 70°C for 20 min. After being stained, the sections were mounted with mounting solution. Slides were imaged by confocal microscopy with an AX microscope (Nikon) (**Fig. 2A, H; 3A; Supplemental Fig. S1A**).

#### Whole-mount staining of meninges attached to the brain

Fresh heads were collected from E11.5 or E12.5 *Cx3cr1-GFP*^+/−^ embryos and fixed in 4% paraformaldehyde (PFA) for 4 h on ice. After washing with PBS, the skin was carefully removed, and the brains were dissected with the meninges left intact. The brains with attached meninges were divided into rostral, middle, and caudal regions, which were processed separately for whole-mount immunostaining. Samples were incubated overnight at 4°C with chicken anti-GFP polyclonal antibody (1:1,000; Aves Labs, Cat# GFP-1020, RRID:AB_10000240), goat anti-CD31 antibody (1:300; R&D Systems, Cat# AF3628, RRID:AB_2161028), and guinea pig anti-laminin antibody (1:1,000; FUJIFILM Wako, Cat# 010-28821, RRID:AB_3754827). After washing with PBS containing 0.01% Triton X-100, samples were incubated with the following secondary antibodies: Alexa Fluor 488-conjugated anti-chicken IgY antibody (1:1,000; Invitrogen, Cat# A78948, RRID:AB_2921070), Alexa Fluor 546-conjugated anti-goat IgG antibody (1:1,000; Invitrogen, Cat# A11056, RRID:AB_2534103), and Alexa Fluor 647-conjugated anti-guinea pig IgG antibody (1:1,000; Jackson ImmunoResearch Laboratories, Cat# 706-606-148, RRID:AB_2340477). After washing, the stained tissue was mounted in Fluoromount-G (SouthernBiotech) between two coverslips separated by spacers made from plastic tape and imaged immediately using an AX confocal microscope (Nikon) with a 40× objective. Z-stacks were acquired at 1-µm intervals (**Fig. 2C, F**).

#### Treatment with TGF-β1 and TGF-β receptor inhibitors

For the experiments shown in **Fig. 4**, TGF-β receptor signaling was inhibited with the TGF-β type I receptor inhibitor SB431542 (5 μM; Selleck, Cat# S1067). SB431542 was added to the culture medium at the start of direct co-culture with NPCs and maintained throughout the 2-day culture period. An equivalent volume of vehicle was added to the control cultures.

For the experiments shown in **Fig. 5**, recombinant mouse TGF-β1 (20 ng/ml; R&D Systems, Cat# 7666-MB-005) was added to cultures of E13.5 or E17.5 meningeal macrophages. An equivalent volume of vehicle was added to the control cultures. Cells were cultured for 2 days and subsequently collected for flow cytometric analysis.

#### Quantitative real time PCR

Total RNA was extracted from CD11b-positive cells isolated from the meninges of E13.5 and E17.5 mouse embryos using a QIAGEN RNA extraction kit. Meningeal cells from four to five embryos were pooled for each sample. Approximately 100 ng of total RNA was reverse-transcribed into first-strand cDNA using SuperScript III reverse transcriptase (ThermoFisher Scientific) in the presence of RNase inhibitor (Thermo Fisher Scientific).

Quantitative real-time PCR was performed using SYBR Green Real Time PCR Master Mix (Toyobo, Cat# QPS-201) on an AriaMx Real-Time PCR System (Agilent). Amplification was followed by melting curve analysis from 60°C to 95°C to confirm amplification specificity. *Hprt* was used as the reference gene because its expression was stable across the developmental stages and experimental samples analyzed. The threshold cycle (Ct) values of target genes were normalized to those of *Hprt*, and relative expression levels were calculated using the ΔΔCt method. The following primer sequences were used: 5’-TGC TCC AAA CCA CAG AGT AGG C-3’ (sense) and 5’- CCC AGA ACA CTA AGC CCA TTG C -3’ (antisense) for *Tgfbr1*; 5’-CCA TCC ATC CAC TGA AAC ATT TTA AC -3’ (sense) and 5’- GAC AGT CTC ACA TCG CAA AAC TTG-3’ (antisense) for *Tgfbr2*; 5’-CTG GTG AAA AGG ACC TCT CGA AG-3’ (sense) and 5’-CCA GTT TCA CTA ATG ACA CAA ACG-3’ (antisense) for *Hprt*.

#### Quantification and statistical analysis

Quantitative data are presented as individual values overlaid on bars representing the mean; error bars indicate SD. Statistical differences between groups were analyzed using R software by the Mann–Whitney U test for two-group comparisons, the Steel–Dwass test for multiple comparisons and Pearson’s chi-squared test for contingency tables evaluating macrophage migration patterns. All statistical tests were two-tailed, and *p* < 0.05 (*) was considered significant (\*\*\**p* < 0.001, \*\**p* < 0.01, \**p* < 0.05, or n.s., not significant). Individual values were plotted as circles in bar graphs. As for histological analyses, the average value of four or six sections from each animal is plotted. The number of samples examined in each analysis is shown in the corresponding figure legend. No randomization was used, and no samples were excluded from the analysis. No statistical methods were used to predetermine the sample size owing to experimental limitations.

## Legends for Supplemental Figures

**Supplemental Figure S1. Developmental distribution of CD206- and P2RY12-positive cells in the embryonic pallium, related to Figure 3**

(**A**) Representative immunohistochemical images of the pallium at E12.5, E14.5, and E16.5 stained for CD206, P2RY12, and Iba1, with DAPI counterstaining. Scale bar, 100 μm. (**B**) Quantification of the proportions of CD206-positive and P2RY12-positive cells among Iba1-positive cells in the developing pallium at E12.5, E14.5, and E16.5. Statistical significance was assessed using the two-sided Steel–Dwass test (*n* = 6 embryos per stage). Data are presented as the mean, with SD shown only in the positive direction.

**Supplemental Figure S2. Flow cytometric analysis of macrophage phenotypes following co-culture with NPCs or CD11b-negative brain cells, related to Figure 3**.

Representative flow cytometry gating strategy used to identify the CX3CR1-positive macrophage population. Cells were sequentially gated based on FSC/SSC, singlets and live cells, followed by selection of the CX3CR1-positive population (P5, shown in green). Representative flow cytometric plots showing CD206 and P2RY12 expression in CX3CR1-positive macrophages cultured alone (Mf-only), co-cultured with NPCs (+NPCs), or co-cultured with unfractionated CD11b-negative brain cells (+CD11b-negative cells).

**Supplemental Figure S3. Flow cytometric analysis of macrophage phenotypes following co-culture with NPCs and ECs, related to Figure 4**.

Representative flow cytometric plots showing CD206 and P2RY12 expression in CX3CR1-positive macrophages cultured alone (Mf-only), with NPCs (+NPCs), with ECs (+ECs), or with both NPCs and ECs (+NPCs/ECs).

**Supplemental Figure S4. Flow cytometric analysis of macrophage phenotypes following co-culture, related to Figure 4**

(**A**) Quantification of CD206-positive, CD206-single-positive, P2RY12-positive, P2RY12-single-positive, CD206/P2RY12-double-positive, and CD206/P2RY12-double-negative populations among total CX3CR1-positive cells under the Mf-only, +NPCs, +ECs, and +NPCs/ECs conditions corresponding to **Fig. 4B and C**. Statistical significance was assessed using the two-sided Steel–Dwass test (*n* = 5 independent experiments). Data are presented as the mean, with SD shown only in the positive direction. (**B**) Quantification of CD206-positive and P2RY12-positive populations among CX3CR1-positive macrophages cultured alone (Mf-only), directly co-cultured with NPCs (NPC-D), indirectly co-cultured with NPCs using a Transwell system (NPC-T), or directly co-cultured with NPCs in the presence of SB431542 (NPC-D/I), corresponding to **Fig. 4D and E**. Statistical significance was assessed using the two-sided Steel–Dwass test (*n* = 5 independent experiments). Data are presented as the mean, with SD shown only in the positive direction.

**Supplemental Figure S5. Effects of cellular proximity and TGF-β receptor signaling on the NPC-induced macrophage phenotypic transition, related to Figure 4**.

Representative flow cytometric plots showing CD206 and P2RY12 expression in CX3CR1-positive macrophages cultured alone (Mf-only), directly co-cultured with NPCs (NPC-D), indirectly co-cultured with NPCs using a Transwell system (NPC-T), or directly co-cultured with NPCs in the presence of SB431542 (NPC-D/I).

**Supplemental Figure S6. Developmental-stage-dependent acquisition of microglial properties by meningeal macrophages, related to Figure 5**.

Representative flow cytometric plots showing CD206 and P2RY12 expression in CX3CR1-positive meningeal macrophages isolated from E13.5 embryos and analyzed under Mf-only, +NPCs, and +TGF-β1 conditions.

**Supplemental Figure S7. Developmental-stage-dependent acquisition of microglial properties by meningeal macrophages, related to Figure 5**.

Representative flow cytometric plots showing CD206 and P2RY12 expression in CX3CR1-positive meningeal macrophages isolated from E17.5 embryos and analyzed under Mf-only, +NPCs, and +TGF-β1 conditions.

## Legends for Supplemental Videos

**Video S1. Macrophage behavior at the meningeal surface, related to Figure 1**.

Time-lapse imaging of organotypic sagittal brain slices prepared from an E11.5 and E12.5 *Cx3cr1-GFP*^+/−^ embryos, showing a CX3CR1-GFP-positive macrophage entering the developing pallium from the meningeal surface. Images were acquired at 5-min intervals for 8 h. Scale bar, 20 μm.

**Video S2. Three-dimensional visualization of a representative macrophage at the pial surface of the rostral pallium at E11.5, related to Figure 2**.

Three-dimensional reconstruction of a whole-mount confocal z-stack from an E11.5 *Cx3cr1-GFP*^+/−^ embryo, focusing on a representative CX3CR1-GFP-positive macrophage in relation to the laminin-positive pial basement membrane and CD31-positive vasculature. The imaged region corresponds to the green square in **Fig. 2A**, with the corresponding XY images shown in **Fig. 2H**. Sequential viewing through the z-stack shows the spatial relationship between the macrophage and a local discontinuity in the pial basement membrane. Scale bar, 20 μm.

## Declaration of generative AI and AI-assisted technologies in the manuscript preparation process

During the preparation of this work, the authors used ChatGPT (OpenAI) to assist with English language editing. The authors reviewed and edited the output as needed and take full responsibility for the content of the published article.

## Notes

### Competing Interest Statement

The authors have declared no competing interest.

