## Supplemental Figures for "Embryonic macrophages chronologically change their brain-entry routes and competence for neural progenitor-induced microglial differentiation"

### Supplemental Figure S1

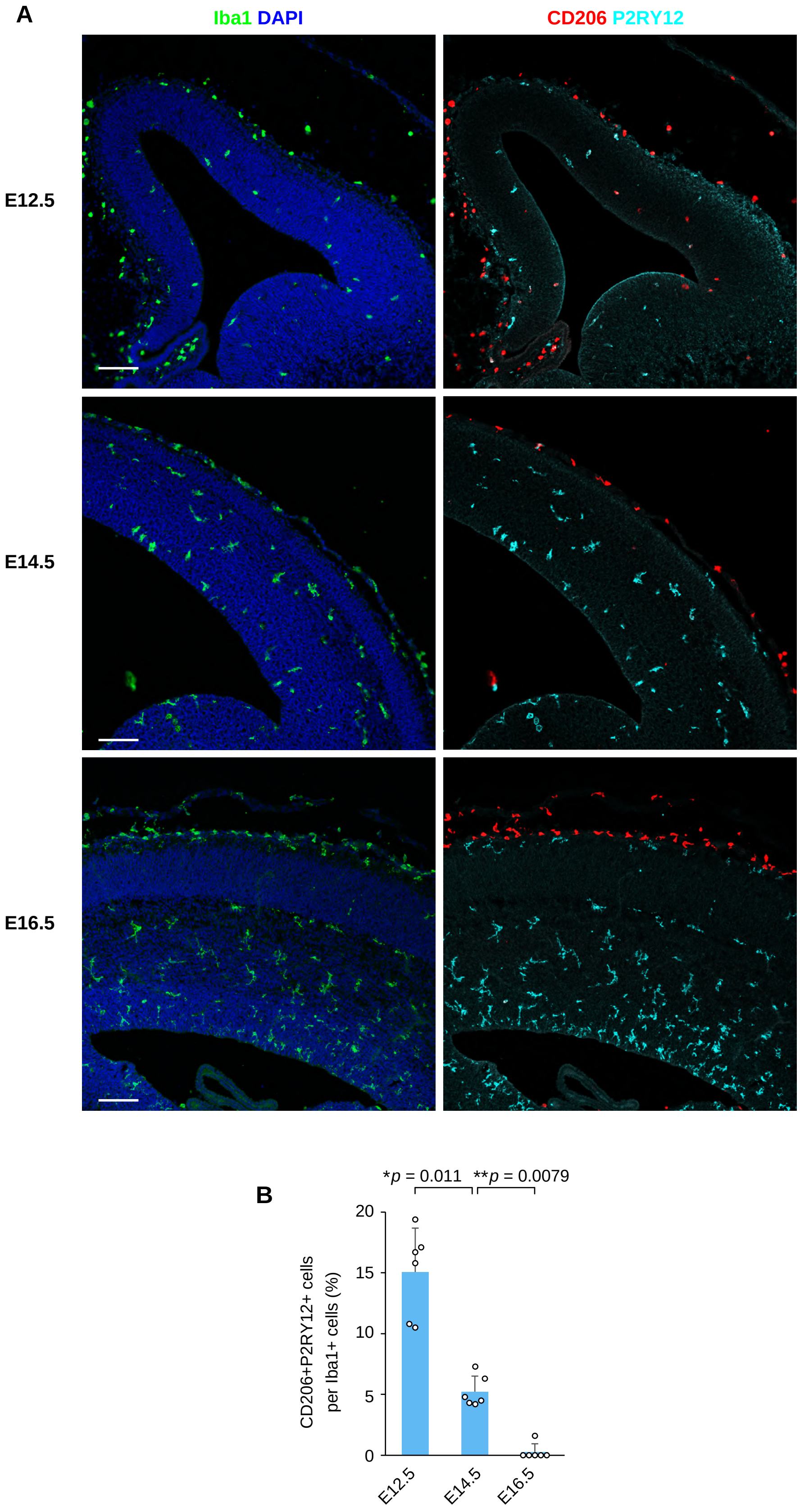

### Supplemental Figure S2

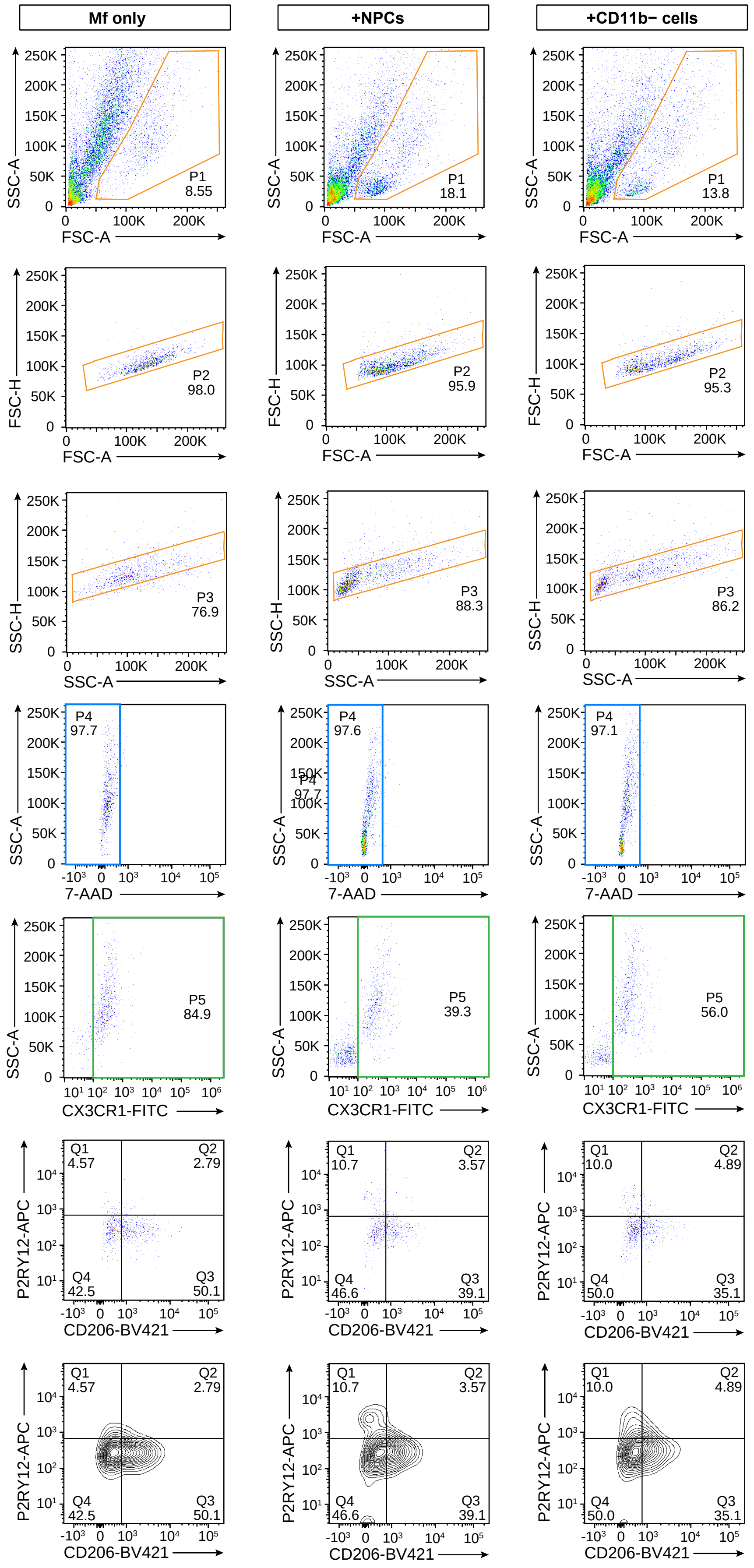

### Supplemental Figure S3

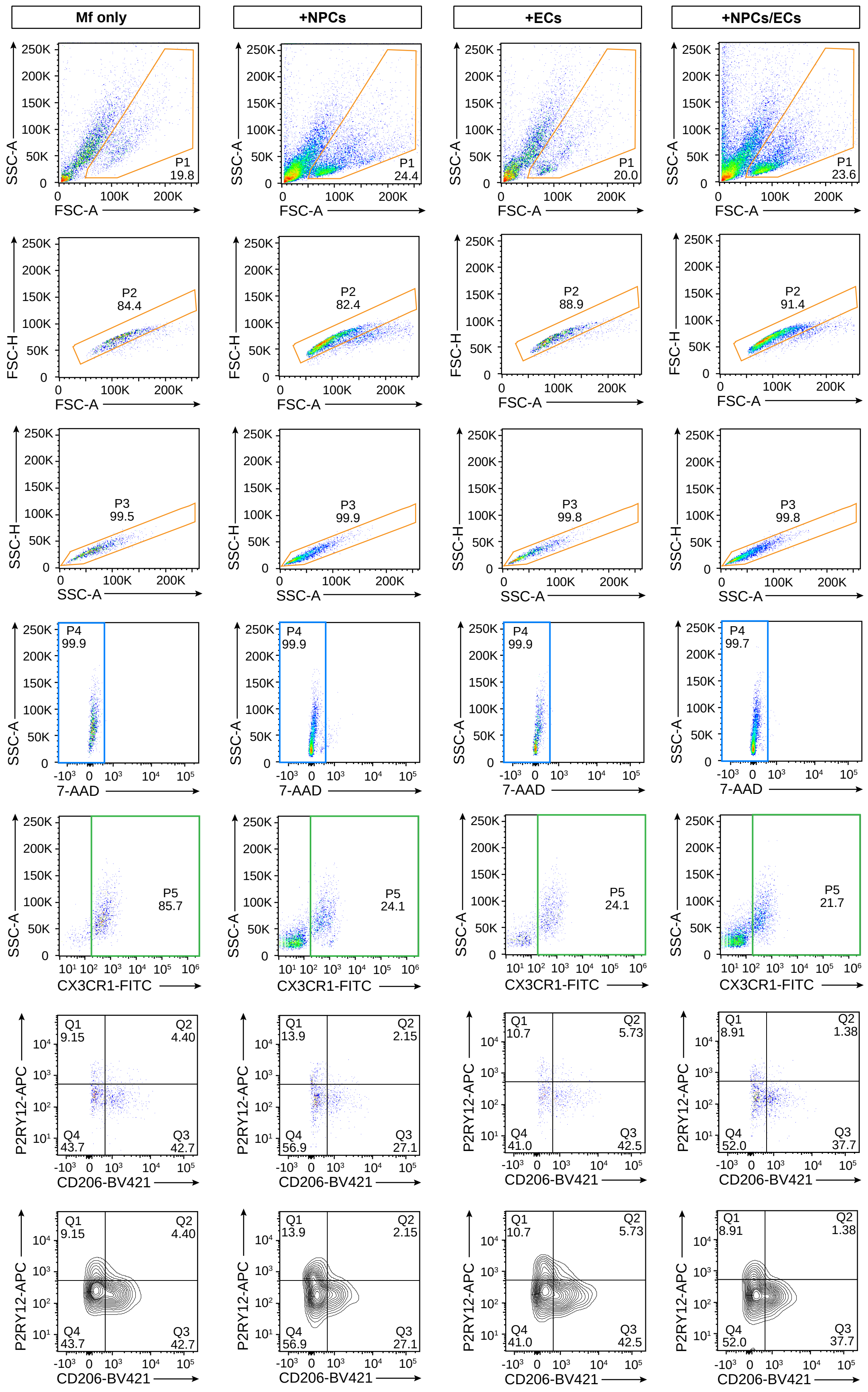

**Supplemental Figure S4**

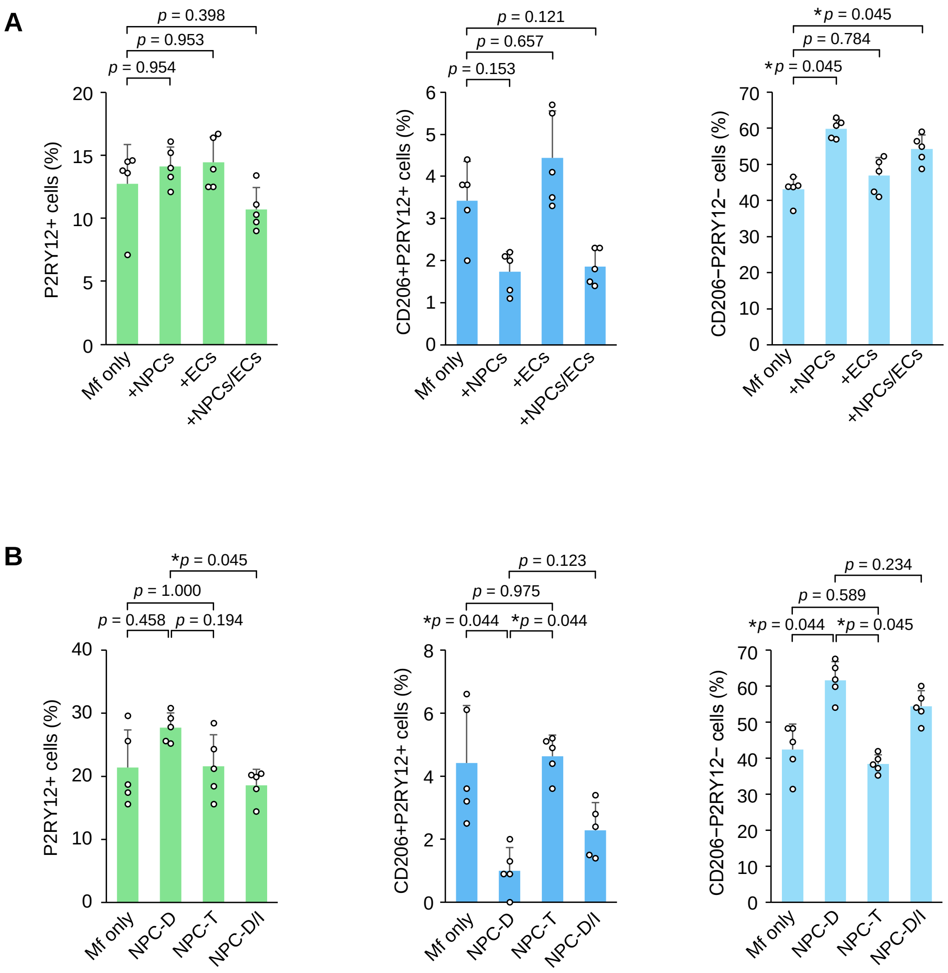

### Supplemental Figure S5

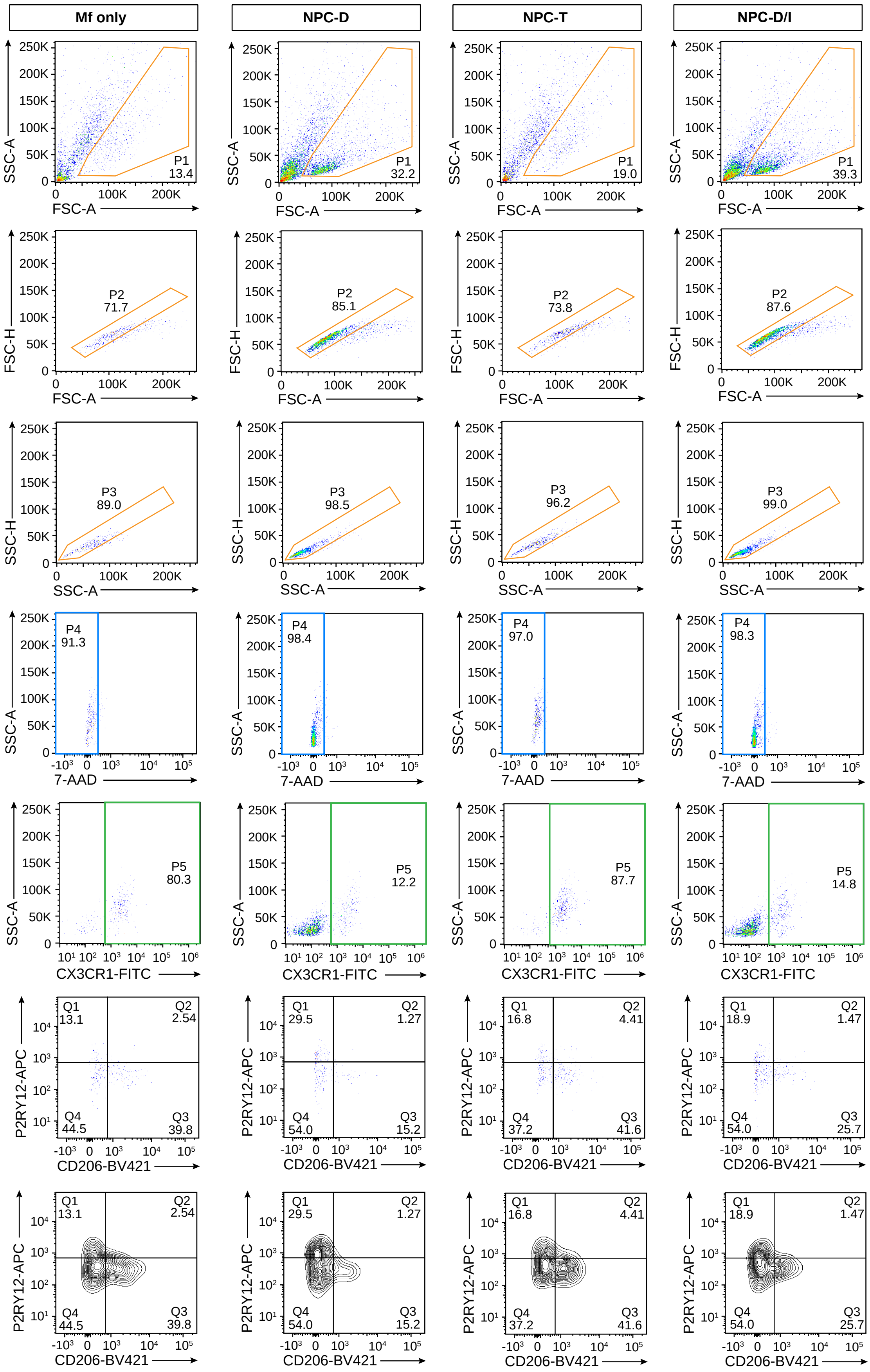

### Supplemental Figure S6

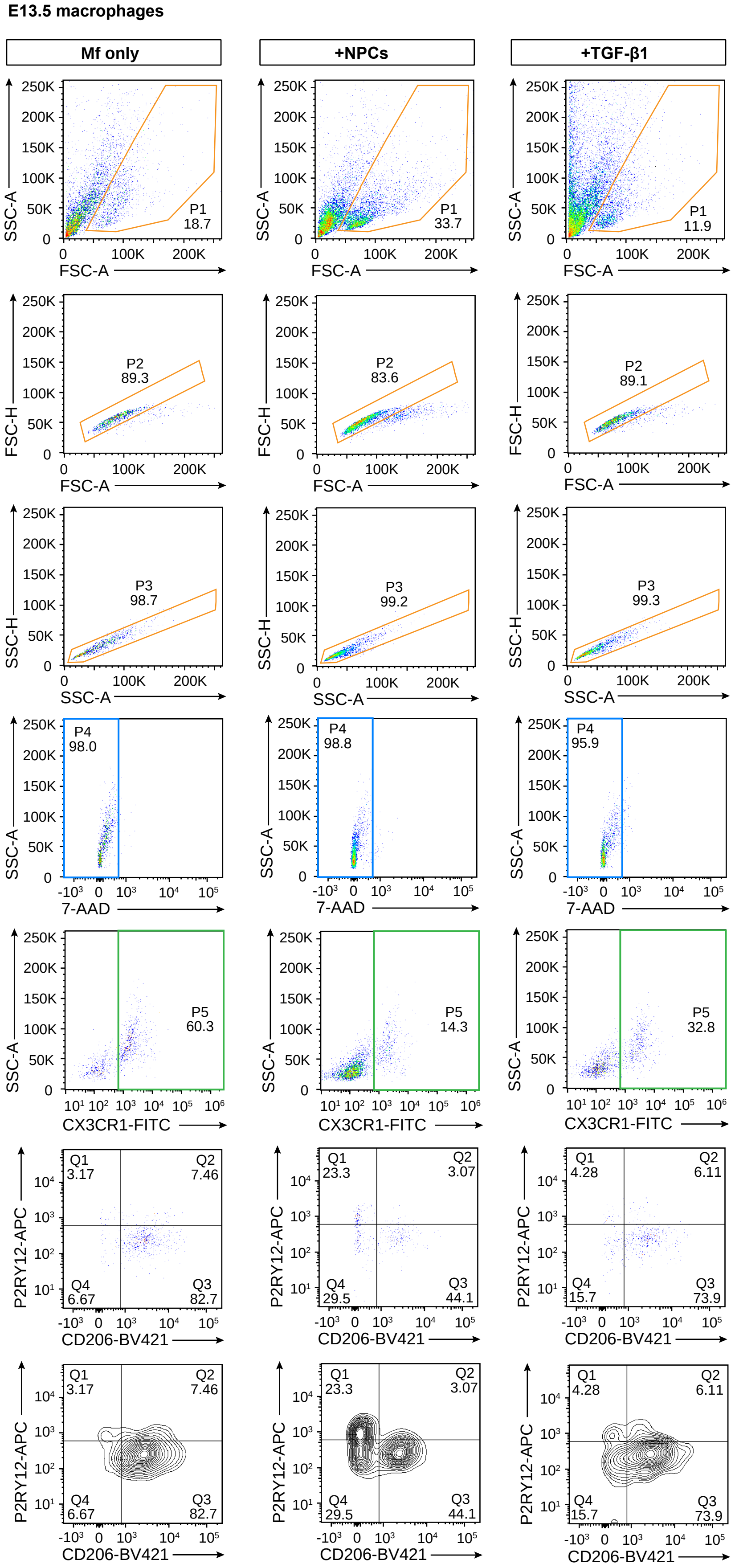

#### Supplemental Figure S7

**E17.5 macrophages**

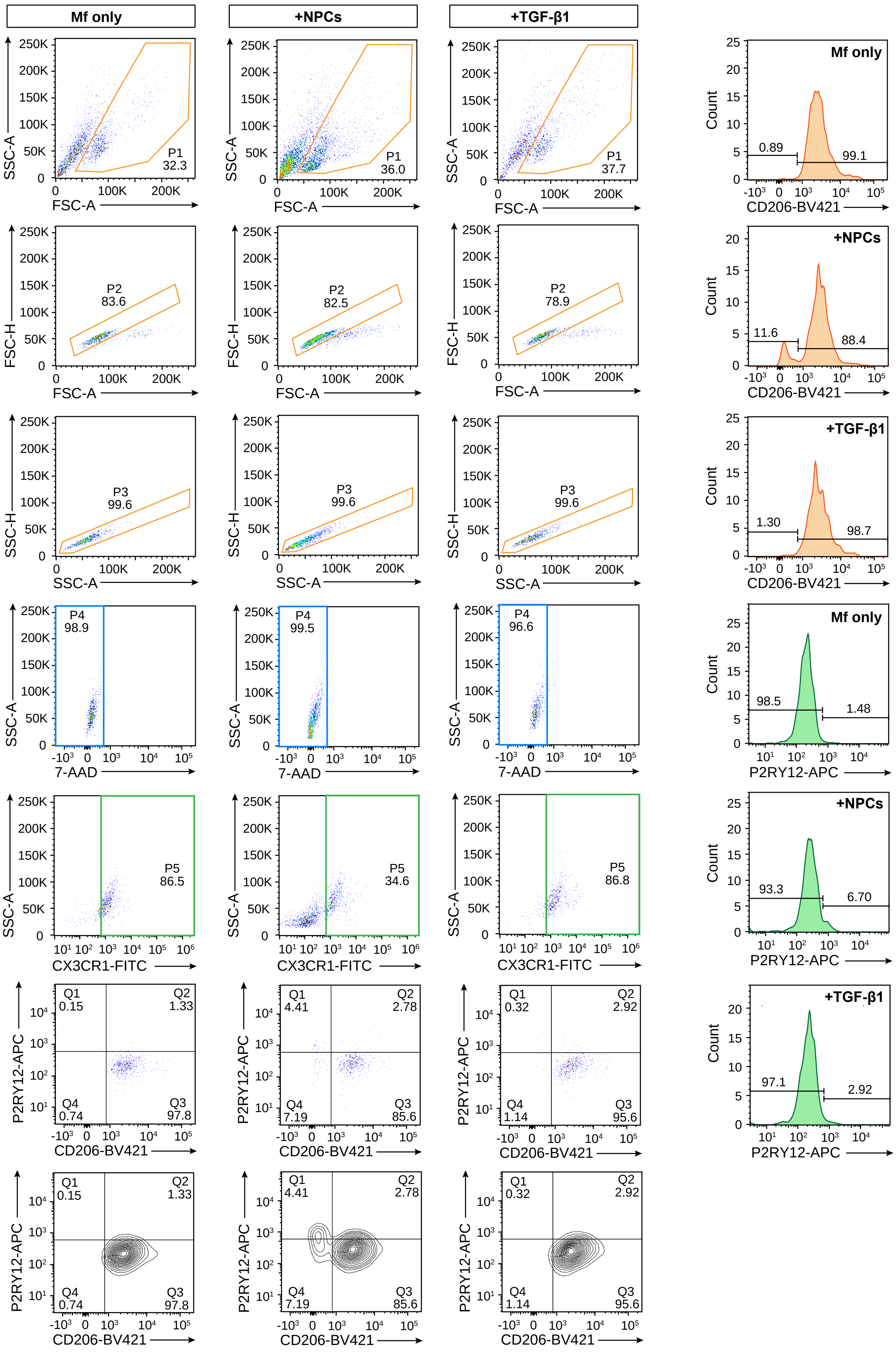
